# Cytotoxic T cells targeting lytic KSHV gene products infiltrate Kaposi sarcoma tumors

**DOI:** 10.64898/2026.06.05.730414

**Authors:** Shashidhar Ravishankar, Andrea M.H. Towlerton, Iyabode L. Tiamiyu, Chris P. Miller, Peter Mooka, Janet Nankoma, James Kafeero, Dennis Mubiru, Semei Sekitene, Lauri D. Aicher, David G. Coffey, Lazarus Okoche, Prisca Atwinirembabazi, Joseph Okonye, Kelvin R. Mubiru, Jessica White, Lichen Jing, David M. Koelle, Warren T. Phipps, Edus H. Warren

**Affiliations:** Translational Science and Therapeutics Division, Fred Hutchinson Cancer Center, Seattle, Washington, United States of America; Hutchinson Centre Research Institute – Uganda, Kampala, Uganda; Department of Laboratory Medicine and Pathology, University of Washington, Seattle, Washington, United States of America; Molecular Medicine & Mechanisms of Disease Graduate Program, University of Washington, Seattle, Washington, United States of America; Uganda Cancer Institute, Kampala, Uganda; Division of Myeloma, Sylvester Comprehensive Cancer Center, University of Miami, Miami, Florida, United States of America; Department of Medicine, University of Washington, Seattle, Washington, United States of America; Vaccine and Infectious Disease Division, Fred Hutchinson Cancer Center, Seattle, Washington, United States of America; Department of Global Health, University of Washington, Seattle, Washington, United States of America; Benaroya Research Institute, Seattle, Washington, United States of America

## Abstract

Kaposi sarcoma-associated herpesvirus (KSHV) is the etiologic agent of Kaposi sarcoma (KS), primary effusion lymphoma (PEL), and KSHV-associated multicentric Castleman’s disease (MCD), malignancies that predominantly arise in the context of T-cell deficiency. Unlike responses to other human herpesviruses such as EBV and CMV, KSHV-specific T-cell responses detected in blood have been described as heterogeneous and low-intensity. Hypothesizing that KSHV-specific T cells are recruited to KS tumors, we analyzed the T-cell receptor (TCR) repertoire of biopsies from 144 Ugandan adults with KS (106 people living with HIV [PLWH], 38 HIV-seronegative) and identified >4,000 αβ TCRs with predicted specificity for KSHV- or HIV-encoded peptides presented by specific MHC alleles. We tested 14 putative KSHV- or HIV-specific TCRs for recognition of cells presenting cognate peptides in the predicted MHC context. Three novel HIV-specific TCRs, found only in tumors from PLWH, exhibited high-avidity, MHC-restricted recognition of HIV *Vpr* and *Nef* peptides previously identified as CD8**^+^** T-cell targets. Four KSHV-specific TCRs, detected in tumors from both PLWH and HIV-seronegative individuals, recognized peptides encoded by the lytic KSHV genes *ORF6*, *ORF57*, and *ORF59*. The *ORF6*- and *ORF57*-specific TCRs were observed in multiple individuals and constitute the first examples of public T-cell responses to KSHV. We then confirmed that these four KSHV-specific TCRs recognized KSHV-infected cells undergoing lytic reactivation. Identification of TCRs specific for KSHV lytic gene products will enable the development of T-cell-based therapies for KS and other KSHV-associated diseases.

**Author summary:** Kaposi sarcoma-associated herpesvirus (KSHV) is the only oncogenic human virus for which there is no effective vaccine, antiviral, or immunotherapeutic strategy that directly targets the virus. KSHV is the causative agent of Kaposi sarcoma and primary effusion lymphoma, cancers that primarily affect immunocompromised individuals, most commonly people living with HIV in sub-Saharan Africa, where KSHV is endemic and HIV infection is prevalent. In this study we defined the antigenic specificity of novel, putative KSHV- and HIV-specific T-cell receptors (TCRs) that are carried in T cells commonly infiltrating KS tumor biopsies. We show that T cells engineered to express KSHV-specific TCRs exhibit high-avidity, MHC-restricted recognition of KSHV-infected cells presenting naturally processed peptides encoded by KSHV lytic genes. Our findings provide a blueprint for dissecting KSHV-specific T-cell immunity and advancing the development of immunotherapeutic strategies for the prevention or treatment of KSHV-associated diseases.

## Introduction

Kaposi sarcoma-associated herpesvirus (KSHV) is the etiologic agent of Kaposi sarcoma (KS), primary effusion lymphoma (PEL), and KSHV-associated multicentric Castleman’s disease (MCD) [1]. KS remains a significant cause of cancer-related morbidity and mortality worldwide, particularly among people living with HIV (PLWH) and in sub-Saharan Africa (SSA), where KSHV is endemic and HIV-1 infection is prevalent [2, 3]. Over 85% of global KS-related deaths occur in SSA, and more than 80% of KS cases are attributable to HIV infection [2, 3]. Management of KSHV-associated diseases in SSA relies primarily on chemotherapy, combined with antiretroviral therapy (ART) for PLWH [4]. Immune checkpoint inhibitors and antiviral agents have shown modest activity in subsets of patients with KS [5–9]. The immunomodulatory agent pomalidomide has been approved by the FDA for the treatment of KS in PLWH or individuals without HIV infection [10], but this drug is not readily available in SSA. To date, no virus-specific therapies have been approved for the treatment of KS, PEL, or KSHV-MCD.

In SSA, KSHV infection primarily occurs in childhood [11–13], while KS and other KSHV-associated diseases typically develop many years, often decades later, most commonly in the setting of T-cell deficiency and dysfunction resulting from chronic HIV infection [14]. The incidence of KS is also significantly increased in recipients of allogeneic solid organ or hematopoietic cell transplantation [15, 16]. These observations suggest that loss or impairment of pre-existing KSHV-specific T-cell immunity underlies the pathogenesis of KS and other KSHV-associated diseases.

Previous studies of asymptomatic KSHV-infected individuals and those with KSHV-associated disease have identified several peptides encoded by KSHV genes that are recognized by CD8**^+^** T cells circulating in blood [17–23]. Focused studies of CD8**^+^** and CD4**^+^** T-cell responses to defined KSHV-encoded peptides have reported that these responses are stronger and quantitatively larger in the blood of asymptomatic KSHV-infected individuals than in individuals with KS [21, 24]. Subsequent studies have taken a variety of experimental approaches to define the breadth and magnitude of the CD8**^+^** and CD4**^+^**T-cell response KSHV in the blood of KSHV-infected PLWH and KSHV-infected, HIV-seronegative individuals [25–32]. Detection and characterization of KSHV-specific T-cell responses in blood, however, has often proven to be challenging, which has been attributed to their heterogeneity, low frequency, and low intensity.

Our group has consequently focused on T cells infiltrating KS tumors to characterize the T-cell response to KSHV. Transcriptional profiling by our group [33] and others [34, 35] shows consistent expression of latent and lytic KSHV genes within KS tumors as well as expression of HIV genes in KS tumors from PLWH. In this study we analyzed KS tumor transcriptomes using CIBERSORTx [36, 37] and found that CD8^+^ T cells and M2-polarized macrophages are the most abundant types of immune cells. We performed adaptive immune receptor repertoire sequencing (AIRR-seq) of 299 KS tumor biopsies and 64 samples of uninvolved axillary skin from 106 PLWH with KS and 38 HIV-seronegative individuals enrolled on a prospective natural history study [38] and found that ∼25% of TCRs detected in KS tumors clustered into antigenic specificity groups predicted to recognize similar or identical peptides with shared MHC restriction. Interrogation of public TCR databases revealed that these TCRs have not previously been associated with T-cell responses to known pathogens or tissue antigens. We hypothesized that the clustered TCRs with undefined antigenic specificity that we observed in KS tumors may be enriched for TCRs specific for HIV or KSHV. We investigated the antigenic specificity of 16 of these clustered TCRs and identified three novel high avidity TCRs recognizing HIV-encoded peptides and four novel high avidity TCRs recognizing peptides encoded by KSHV lytic genes. T cells engineered to express the KSHV-specific TCRs showed MHC-restricted recognition of KSHV-infected cells that had been treated to trigger lytic replication of KSHV.

## Results

### KSHV and HIV gene expression in KS tumors

Our group previously reported that RNA-sequencing (RNA-seq) of 39 KS tumors obtained from PLWH (termed epidemic KS) detected expression of both latent and lytic KSHV genes within the tumor microenvironment (TME) [33]. We performed RNA-seq on 12 additional KS tumors from adults without HIV infection (termed endemic KS) and similarly identified consistent expression of latent and lytic cycle genes (**Fig 1**). To extend these findings, we leveraged published RNA-seq datasets and applied the same analysis to quantify KSHV and HIV viral gene expression. These datasets comprised 22 epidemic KS lesions with paired lesion-adjacent skin, 6 endemic KS lesions with paired lesion-adjacent skin, and 3 non-KS control skin samples [34, 35].

**Figure 1.**
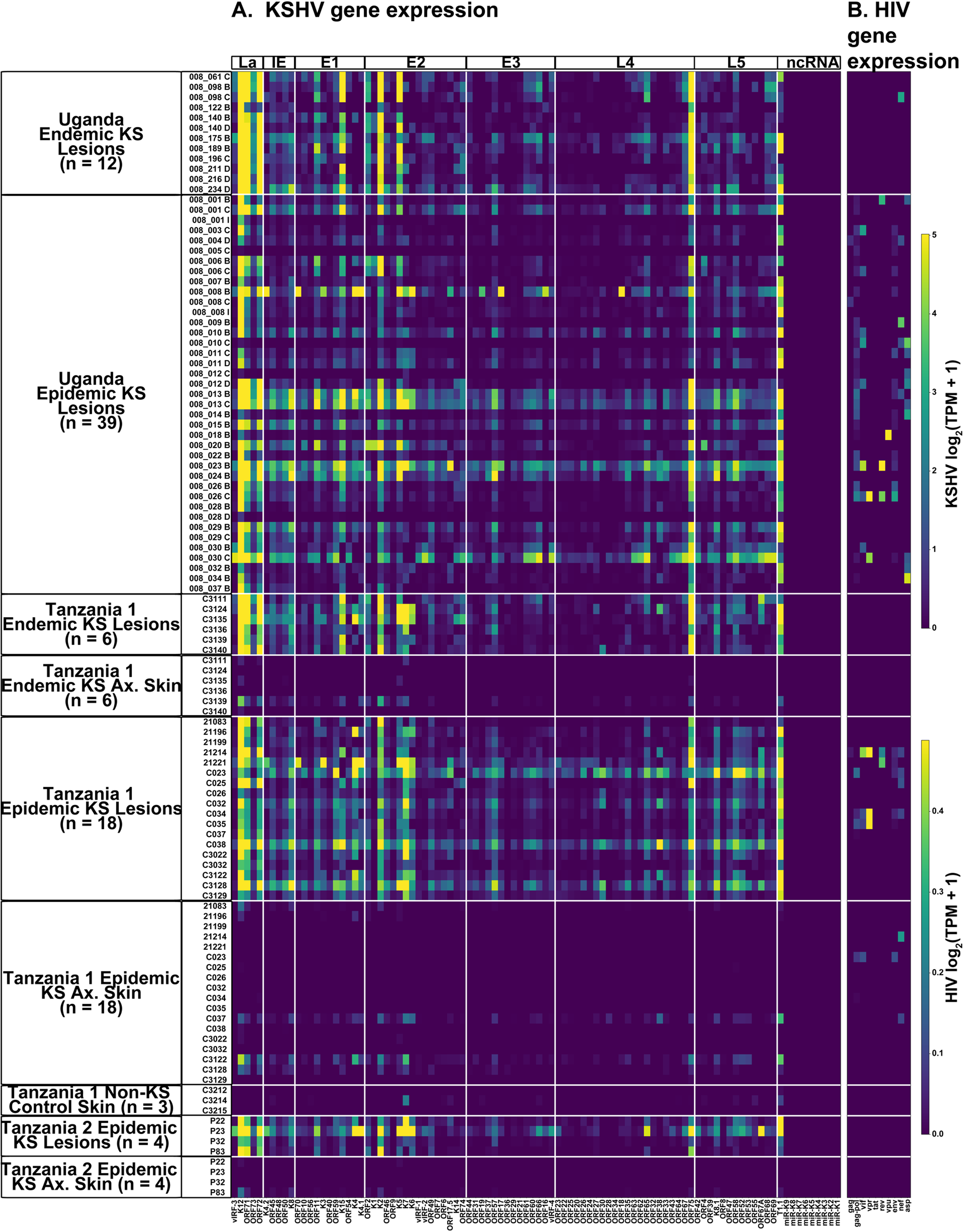
RNA-seq confirms expression of KSHV and HIV genes in KS tumors. (A) Heatmap of KSHV gene expression [log₂(TPM+1)] in 18 endemic KS tumors, 61 epidemic KS tumors, 28 paired axillary skin samples, and 3 non-KS control skin samples, compiled from this and published studies [33–35]. KSHV genes are grouped by kinetic class: latent (La), immediate early (IE), early lytic stages 1, 2, and 3 (E1, E2, E3), late lytic stages 4 and 5 (L4, L5), and non-coding RNAs (ncRNA). (B) Heatmap of HIV gene expression [log₂(TPM+1)] across the same 79 KS tumors, 28 paired axillary skin samples, and 3 non-KS control skin samples; expression is detected predominantly in lesions from people living with HIV (epidemic KS).

Kaposin (*K12*) was the most highly expressed KSHV protein-coding gene in all but 9 of the 79 KS tumor samples (median 107.5 TPM, range 0.1-1644.5 TPM) and was also the most highly expressed gene in the 22 tumors evaluated with a targeted expression assay (**S1 Fig**). Notably, non-coding KSHV RNAs overlapping with the *K7* locus (*PAN/T1.1*) also ranked among the most abundant viral transcripts and exceeded K12 levels in several tumors. Other latency genes, including *ORF71* (median 10.9 TPM) and *ORF72* (median 10.6 TPM), were detected in 76 of 79 tumors, and endemic KS tumors showed 6- to 9-fold higher median transcript levels across genes of the KSHV latency program than epidemic KS tumors. The primary latency gene *ORF73* (LANA-1) was detected at lower transcript levels than other latency genes (median 0.5 TPM) but was present in 76 of 79 KS lesions. The key lytic gene *ORF75* (median 13.3 TPM) ranked third among all KSHV protein-coding genes and was detected in all 79 tumors, reflecting the dynamic state of KSHV within the KS TME. Lytic genes spanning immediate-early (*e.g., K8)*, early (*e.g., K15, K2)* and late (*e.g., K8.1)* stages of viral replication cycle were broadly expressed, with *K2* detected in 76 of 79 and *K15* in 72 of 79 lesions, underscoring the coexistence of latent and lytic KSHV transcriptional programs within the TME. HIV gene expression was detected in 36 of 57 tumors from individuals with epidemic KS (**Fig 1B**). HIV *Gag/Pol* transcripts were detected in 32 of these tumors, indicating active HIV replication within the TME.

### M2 macrophages and CD8^+^ T-cells are the dominant immune cell types in KS tumors

Deconvolution of the RNA-seq data from the 51 KS tumors using CIBERSORTx [36, 37] was performed to estimate the relative size of immune cell subsets in the KS TME. Macrophages with M2 polarization were predicted to be the most common immune cell type, accounting for a mean of 38% of the immune cells in endemic KS tumors and 22% of the immune cells in epidemic KS tumors (**S2A and S2B Figs**). CD8**^+^** T cells were predicted to be the second largest group of immune cells within the TME, accounting for a mean of 18% and 21% of the cells in the endemic and epidemic tumors, respectively. CD4**^+^** resting memory T cells and CD4**^+^** activated memory T cells were predicted to make up 5% and 4%, respectively, of the immune cells in endemic KS tumors but only 1.3% and 0.2% of the immune cells in epidemic KS tumors (**S2B Fig**). Cell type deconvolution of previously published RNA-seq datasets [34, 35] from 28 KS tumor – normal adjacent tissue (NAT) pairs and skin from 3 non-KS control subjects similarly revealed that M2-polarized macrophages and CD8**^+^** T cells were the most common immune cells in the KS tumors (**S2A Fig**). In contrast, deconvolution analysis of RNA-seq data from normal sun-exposed and non-sun-exposed skin from the GTEx consortium [39] showed a predominance of resting mast cells but low levels of M2 macrophages and few, if any, CD8**^+^** T cells (**S2A Fig**).

### KS TIL carry the signature of a polyclonal response to unknown antigens

The consistent expression of KSHV genes in all KS tumors and of HIV genes in KS tumors from PLWH suggested that T cells specific for antigens encoded by KSHV or HIV would be attracted to KS tumors. We performed adaptive immune repertoire sequencing (AIRR-seq) of the TCR repertoire of the tumor-infiltrating lymphocytes (TIL) in 299 KS tumor biopsies obtained from 144 Ugandan adults [38], 106 of whom were PLWH and 38 who did not have HIV infection (**Fig 2A**). We also performed AIRR-seq on biopsies of uninvolved axillary skin from 64 of the 144 individuals. Using standard bulk TCR α and β chain sequencing, we identified a total of 1.4 million TCR α chain sequences and 2.85 million TCR β chain sequences in the 299 KS tumor biopsies. We also performed single-cell RNA sequencing with T-cell receptor VDJ sequencing (scRNAseq + VDJ) on 20 peripheral blood mononuclear cell (PBMC) samples from a subset of 9 individuals (5 PLWH and 4 HIV-seronegative) to enable pairing of TCR α and β chain sequences from those individuals (single-cell VDJ sequencing results provided in supplementary file S1). This effort identified 61,763 T cells that collectively carried 52,590 unique αβ TCRs. High-resolution class I and class II MHC genotyping data were also acquired for all 144 individuals.

**Figure 2.**
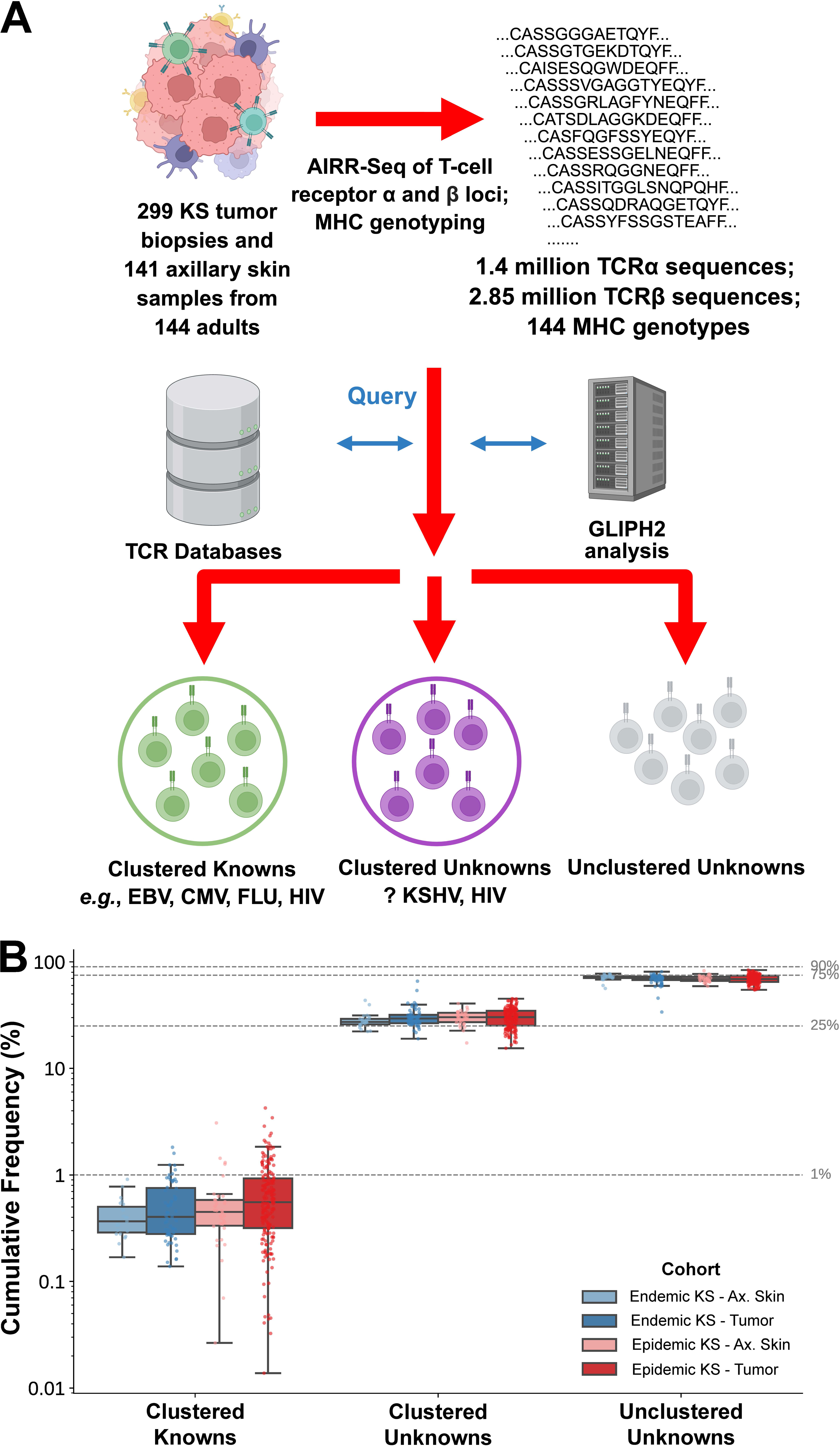
Clusters of T cells with predicted shared specificity for unknown antigens comprise ∼25% of the T-cell repertoire in KS tumors. (A) Schematic of the analytic pipeline. AIRR-seq of the TCRα and TCRβ loci was performed on 299 KS tumor biopsies and 64 axillary skin samples from 144 adults, yielding ∼1.4 million TCRα and ∼2.85 million TCRβ sequences together with paired MHC genotypes. TCRβ sequences were queried against public TCR databases (VDJdb, McPAS-TCR) and clustered by GLIPH2, partitioning the repertoire into three subsets: *Clustered Knowns* - sequences clustering with TCRs of known specificity (e.g., EBV, CMV, influenza, HIV); *Clustered Unknowns* - sequences clustering with one another but not matching any known specificity (candidate KSHV- and HIV-specific TCRs); and *Unclustered Unknowns* - sequences with no cluster assignment. (B) Cumulative frequency of TCRβ sequences assigned to each of the three subsets in axillary skin and tumor biopsies from representative individuals with endemic KS (blue) and epidemic KS (red). Boxes show the median and interquartile range; whiskers extend to 1.5 × IQR; individual samples are overlaid as points. Across all four groups, Clustered Knowns account for <1% of the repertoire, Clustered Unknowns for ∼25%, and Unclustered Unknowns for ∼75%, indicating that a substantial fraction of the KS-infiltrating T-cell repertoire is shared across individuals but not attributable to previously characterized antigenic specificities.

The TCR α and β chain sequences identified in the 299 KS tumor biopsies were compared with the VDJdb [40–42] and McPAS-TCR [43] databases of annotated TCR sequences to identify sequences that have been associated with human T-cell responses to specific microbial or tissue antigens or specific pathological conditions (**Fig 2A**). This identified a small number of sequences, <1% of the total, that have previously been associated with T-cell responses to pathogens such as CMV, EBV, influenza A, and HIV which we have termed clustered knowns (**Fig 2B**). The TCR β chain with CDR3 amino acid sequence CASSVDKGGTDTQYF, for example, was observed in 24 tumors from 13 individuals, all of whom were PLWH and HLA-B*42:01**^+^**, and this sequence has been associated with an HLA-B*42:01-restricted CD8**^+^** T-cell response to the HIV Pol_982-990_ peptide (IIKDYGKQM) [44]. Cells carrying that TCR β chain sequence paired with a TCR α chain with CDR3 sequence CAVSGGGFQKLVF were identified among the T cells whose TCR α and β chains had been captured by scRNAseq. Primary CD8**^+^**T cells engineered to express the CAVSGGGFQKLVF_CASSVDKGGTDTQYF TCR via lentiviral transduction showed reproducible, high avidity recognition (half-maximal recognition, 591 pM) of the IIKDYGKQM peptide when pulsed onto HLA-B*42:01**^+^** target cells. Similarly, the TCR β chain CDR3 sequence CASSLWAGGSNEQFF was observed in 6 KS tumors from 4 PLWH, all of whom were HLA-B*42:01**^+^**, and this sequence has been associated with an HLA-B*42:01-restricted CD8**^+^**T-cell response to the HIV Vpr_34-42_ peptide (FPRPWLHGL) [44]. Cells carrying this TCR β chain paired with a TCR α chain with CDR3 sequence CAPDRAGGTSYGKLTF were also identified in the scRNAseq dataset. Primary CD8**^+^** T cells engineered to express the CAPDRAGGTSYGKLTF_CASSLWAGGSNEQFF TCR via lentiviral transduction likewise showed reproducible, high avidity recognition (half-maximal recognition, 582 pM) of the FPRPWLHGL peptide when pulsed onto HLA-B*42:01**^+^**, but not HLA-B*42:01**^−^**, target cells.

The vast majority (99%) of sequences observed in the TCR repertoires of KS TIL, however, were not listed in VDJdb or the McPAS-TCR database and thus could not be readily associated with previously defined T-cell responses to specific pathogen or tissue antigens. Analysis of the TCR α and β chain datasets with GLIPH2 [45] determined that 24% of the TCRs in KS tumors can be grouped into clusters with predicted similar or identical antigenic specificity, which we have termed “Clustered Unknowns” (**Fig 2A, 2B**). These antigenic specificity clusters show strong association with specific MHC alleles, suggesting polyclonal T-cell responses to MHC-restricted antigens. AIRR-seq of non-contiguous KS tumor biopsies acquired from individual participants before initiation of KS treatment and at subsequent timepoints over the course of 1 year revealed many examples of clustered unknown TCR sequences that could be reliably detected in all tumors acquired over the course of the year, suggesting a persistent T-cell response to the target antigens (**Fig 3**). We hypothesized that many of the TCR sequences that (i) fall into antigenic specificity clusters with high-confidence MHC associations, (ii) are found in multiple non-contiguous KS tumors, and (iii) persist over time, are specific for either KSHV- or HIV-encoded antigens. We further hypothesized that clustered unknown TCRs that are observed only in PLWH are possibly specific for HIV, and that clustered unknown TCRs observed in both PLWH and HIV-seronegative individuals are possibly specific for KSHV.

**Figure 3:**
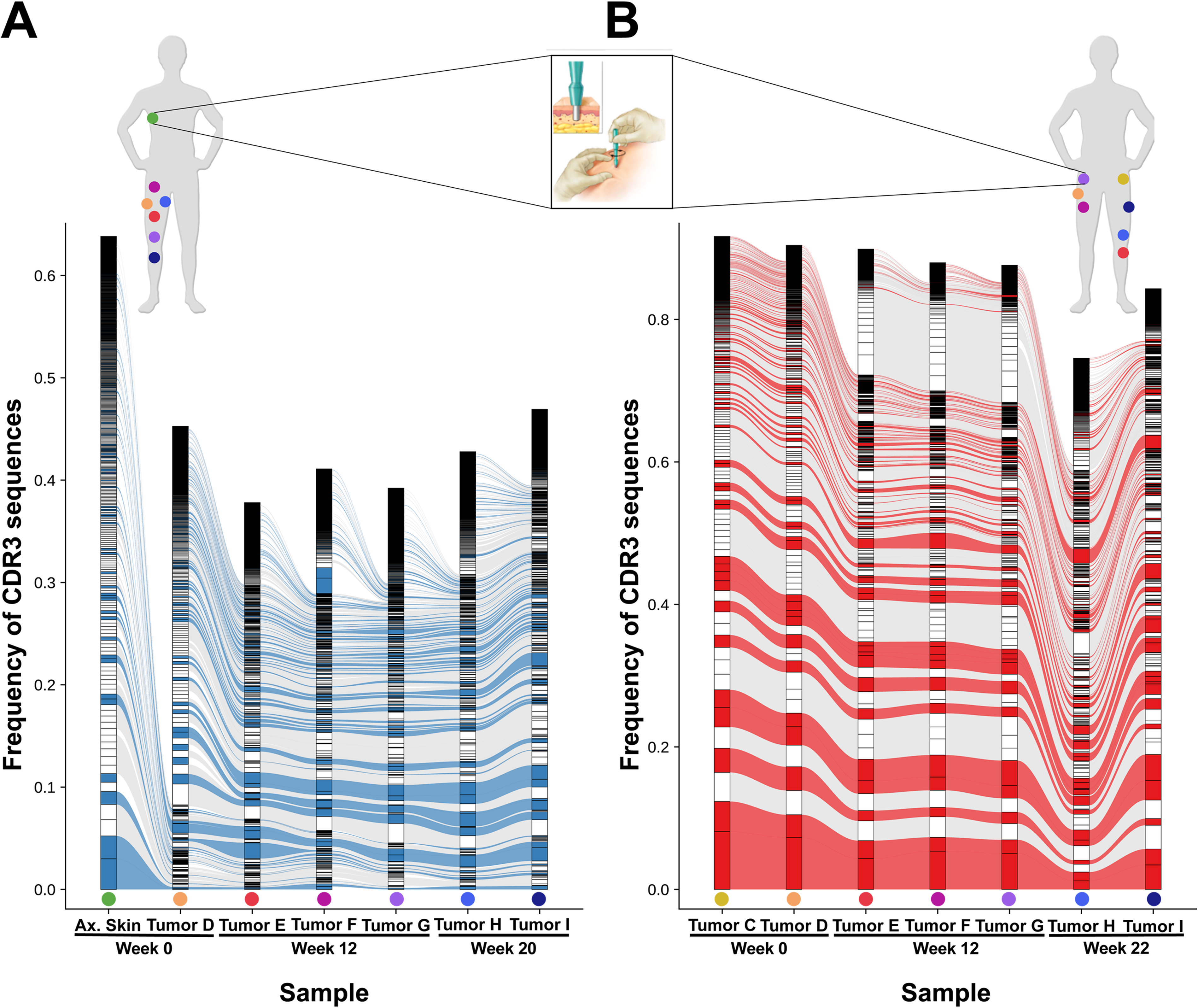
T cells from GLIPH2-defined clusters with specificity for unknown antigens persist in the KS TME across time and space in individuals with endemic and epidemic KS. (A, B) Alluvial plots of the 500 most frequent T-cell receptor β chain sequences observed in tumor samples obtained from multiple anatomic sites and at different time points in two representative individuals with (A) endemic KS (6 tumor biopsies and 1 axillary skin biopsy) and (B) epidemic KS (7 tumor biopsies). Highlighted are persistent TCR β chain CDR3 sequences from GLIPH2-defined “clustered unknown” antigenic specificity groups in endemic (blue) or epidemic (red) KS lesions. Colored dots on the body figures above each alluvial plot denote the sites from which each tissue sample was collected and are reproduced below the corresponding column containing the repertoire data for that tissue sample.

### T cells carrying clustered unknown TCRs seen only in KS tumors from PLWH recognize HIV-encoded peptides

Of the 52,590 unique αβ TCRs captured by scRNAseq, 4,283 were classified as clustered unknowns and were therefore good candidates for KSHV- or HIV-specific TCRs. This subset included many TCRs that were strongly associated with specific class I MHC alleles and observed in KS tumors obtained at multiple time points from PLWH but not observed in tumors from individuals who did not have HIV infection, suggesting that these TCRs might be specific for antigens encoded by HIV. We investigated the antigenic specificity of three clustered unknown TCRs (which we designated TCR-4, TCR-6, and TCR-8**; S1 Table**) that were strongly associated with HLA-B*42:01 and only observed in HLA-B*42:01**^+^** PLWH. TCR-6 is a public TCR that was observed in nine KS tumors obtained from five HLA-B*42:01**^+^** PLWH. TCR-8 is a public TCR that was observed in a distinct set of nine KS tumors obtained from four HLA-B*42:01**^+^**PLWH, while TCR-4 is a private TCR observed in a single HLA-B*42:01**^+^**PLWH.

These TCRs were expressed in Jurkat reporter T cells in which transcriptional activity of the immediate early gene NR4A1 (Nur77) that is induced upon TCR signaling is reported by mNeonGreen fluorescence [46]. The transduced Jurkat reporter T cells were tested for recognition of Epstein Barr Virus-transformed lymphoblastoid cell lines (LCLs) pulsed with 11 HIV-encoded peptides that had previously been shown to be epitopes for HIV-specific, CD8**^+^** HLA-B*42:01-restricted T cells (**S2 Table**) [47]). TCR-6 recognized the HIV Vpr-_34-42_ peptide when presented by HLA-B*42:01**^+^** but not LCLs expressing the irrelevant HLA-B*45:01 (**Fig 4A**). TCR-4 and TCR-8, which our GLIPH2 analysis suggested had similar or identical antigenic specificity, recognized HLA-B*42:01**^+^** but not HLA-B*45:01^+^ LCLs pulsed with the HIV Nef_71-79_ peptide. When expressed in primary human CD8^+^ T cells, all three TCRs were highly avid, with half-maximal specific lysis of peptide-pulsed LCLs at peptide concentrations of ≤5 nM (**Fig 4B**). These experiments identified three novel TCRs recognizing HIV peptides and confirmed the utility of our GLIPH-based prediction of clustered unknowns to prioritize TCRs for investigation.

**Figure 4.**
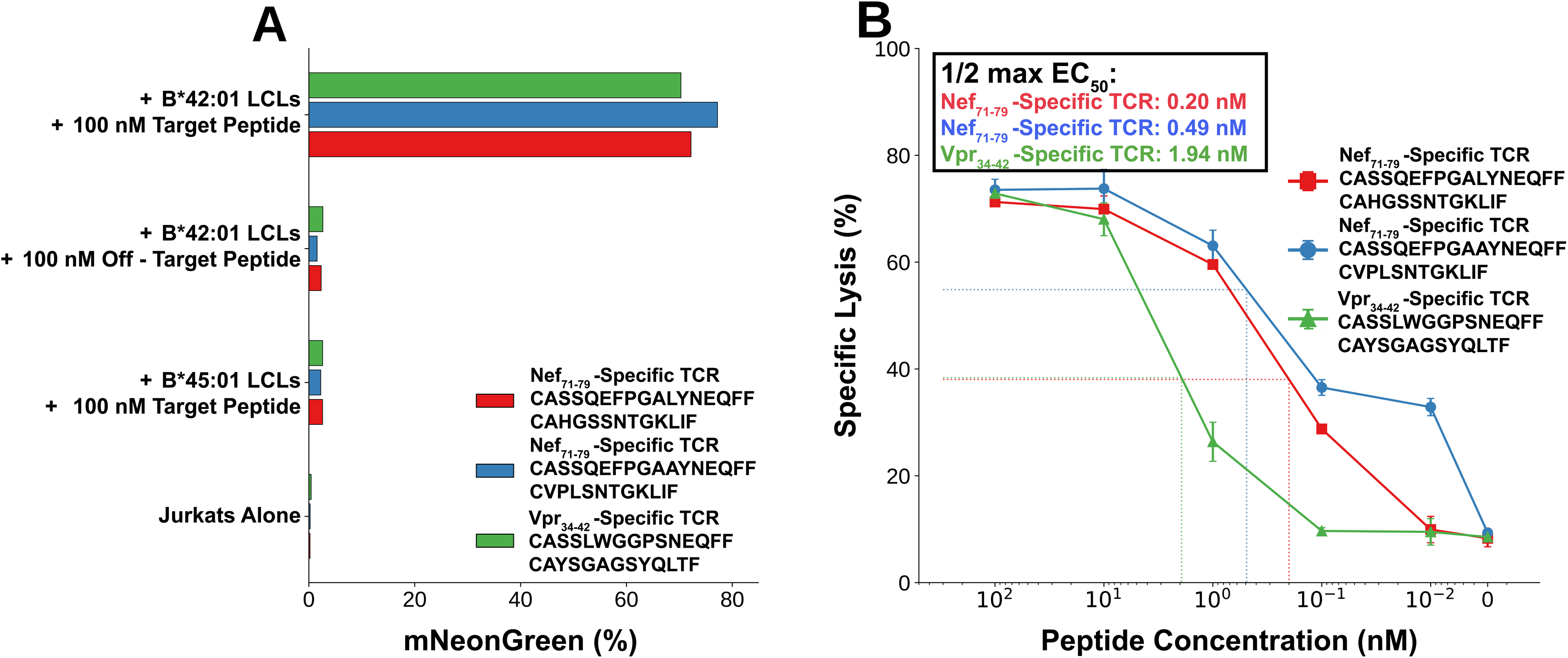
T cells carrying HIV-specific TCRs are observed in KS tumors from PLWH. (A) TCR signaling in Jurkat reporter cells (mNeonGreen fluorescence) transduced with HIV-specific TCRs and co-cultured with LCLs expressing the relevant HLA (B*42:01) or irrelevant HLA (B*45:01) and pulsed with the indicated HIV peptides. (B) Cytotoxicity of HIV-specific TCR-transduced primary human CD8^+^ T cells against HLA-B*42:01^+^ LCL target cells pulsed with the indicated corresponding HIV peptides in a four-hour ^51^Cr release assay at an E:T ratio of 10:1. Mean +/- SEM. Red, TCR-4; Blue, TCR-8; Green, TCR-6.

### T cells expressing public, high-avidity KSHV-specific TCRs infiltrate KS tumors from PLWH and HIV-seronegative individuals

A large subset of clustered unknown TCRs with strong association to specific class I MHC alleles were observed in both endemic and epidemic KS tumors, suggesting the possibility that they recognized KSHV-encoded antigens. A TCR β chain sequence carried in a clustered unknown TCR, which we termed TCR-3 (**S1 Table**), was identified in 29 KS tumors from 8 PLWH and 6 HIV-seronegative individuals, all of whom expressed HLA-B*45:01. T cells carrying this public TCR β chain paired with a specific TCR α chain were captured in the scRNAseq data. Jurkat reporter T cells expressing the αβ TCR-3 were tested for recognition of COS-7 cells transiently co-transfected with plasmids encoding HLA-B*45:01 and pools of plasmids collectively encoding the entire KSHV ORFeome. Screening of iteratively smaller KSHV ORF pools and then minigenes of the KSHV lytic gene ORF6 spanning progressively smaller intervals demonstrated HLA-B*45:01-dependent recognition of a minigene encoding ORF6_329-337_ (**S3 A, B Fig 5A**). Primary CD8**^+^** T cells transduced with TCR-3 demonstrated HLA-B*45:01-dependent cytotoxicity of LCLs pulsed with the ORF6_329-337_ peptide, AEQALHIGA, with half maximal recognition observed at low nanomolar peptide concentrations (**Fig 5A**).

**Figure 5.**
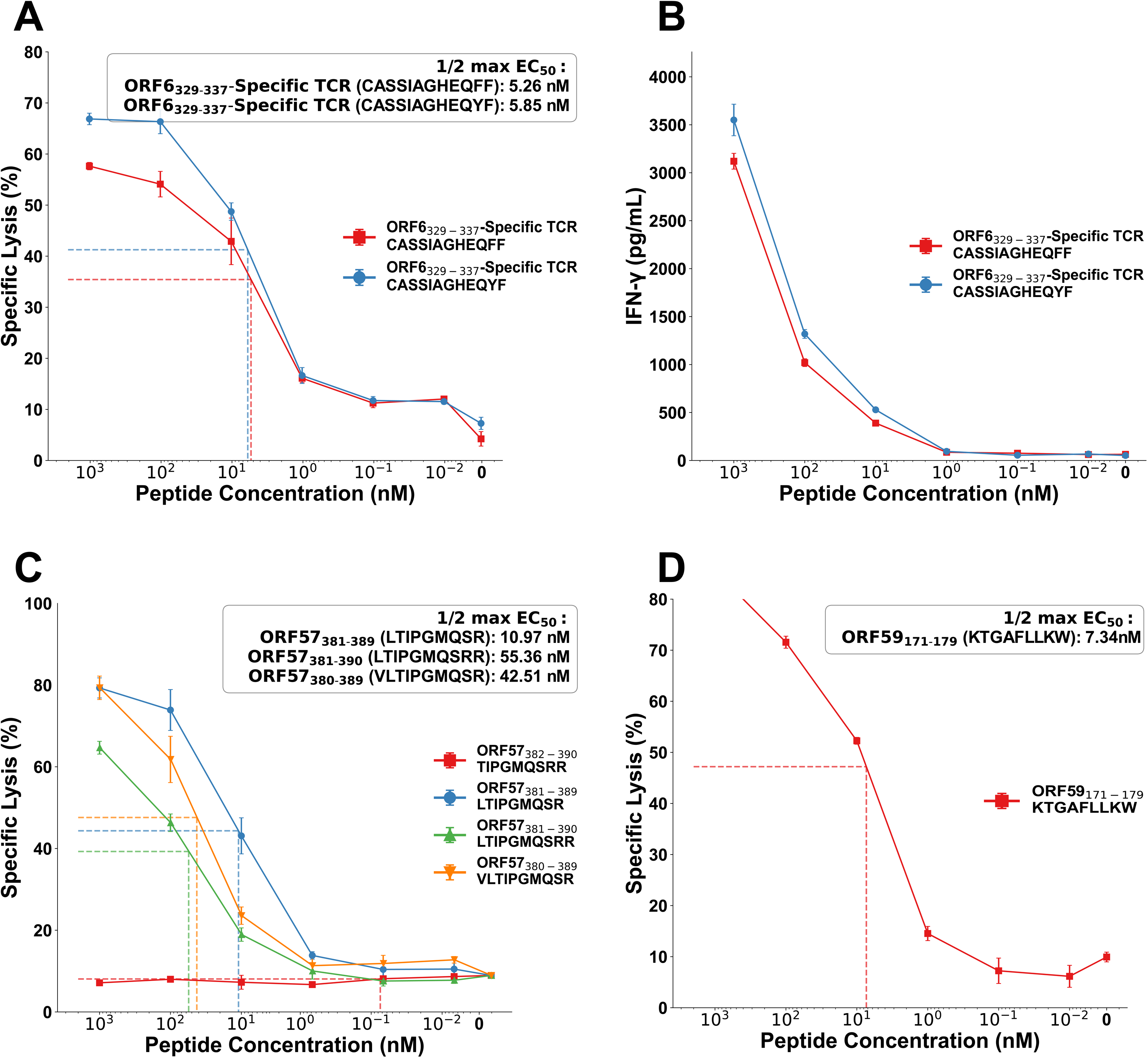
T cells in KS TIL carry high-avidity TCRs specific for KSHV-encoded peptides. **(A)** Cytotoxicity of ORF6-specific TCR-2 (red) or -3 (blue)-transduced primary C8**^+^** T cells against HLA-B*45:01**^+^** LCLs target cells pulsed with the ORF6_329-337_ peptide in a four-hour ^51^Cr release assay at an E:T ratio of 10:1 and **(B)** IFN-γ production. **(C)** TCR signaling in Jurkat reporter cells (mNeonGreen fluorescence) transduced with ORF57-specific TCR-21 and co-cultured with LCLs pulsed with the indicated ORF57 peptides or in **(D)** ORF59-specific TCR-24 and co-cultured with LCLs pulsed with the indicated peptide. Mean +/- SEM.

TCR-3 belongs to an HLA-B*45:01-associated clustered unknown antigenic specificity group that includes a highly homologous TCR, which we termed TCR-2 (**S1 Table**), carrying similar TCR α and β chains with CDR3 sequences mismatched with those of TCR-3 at only one residue each. Like TCR-3, the public TCR β chain carried in TCR-2 was identified in 27 KS tumors from 8 PLWH and 5 HIV-seronegative individuals, all of whom expressed HLA-B*45:01. TCR-2 exhibited prominent intratumoral expansion in two non-contiguous tumors from an HLA- B*45:01**^+^**, HIV-seronegative individual with KS who achieved a complete remission with chemotherapy (**S4A Fig**). TCR-2-expressing Jurkat reporter cells recognized ORF6- and HLA-B*45:01- expressing COS-7 cells and the same panel of progressively smaller minigenes as TCR-3 (**S3C Fig**). TCR-2-expressing primary CD8**^+^** T cells also showed high-avidity, HLA-B*45:01-dependent cytotoxicity against LCLs pulsed with the ORF6_329-337_ peptide, AEQALHIGA, with half maximal recognition observed at low nanomolar peptide concentrations (**Fig 5A**). Both TCR-2 and -3-expressing primary CD8^+^ T cells exhibited robust interferon-γ (IFN-γ) release upon recognition of peptide-pulsed LCLs (**Fig 5B**).

We next investigated a TCR (which we termed TCR-21; **S1 Table**) that demonstrated massive intratumoral expansion in two non-contiguous KS tumors from the same HIV-seronegative individual in which more modest expansion of TCR-2 was observed (**S4A Fig**). We used the COS-7 KSHV ORFeome transfection approach to demonstrate that TCR-21-transduced Jurkat reporter T cells recognized a peptide encoded by the KSHV lytic gene ORF57 and presented by HLA-A*66:01. Evaluation of the ORF57 protein sequence with NetMHCpan [48] identified a nested set of peptide sequences within ORF57: ORF57_381-390,_ ORF57_380-389_, and ORF57_381-389_ – that were predicted to bind to HLA-A*66:01 with high affinity. Primary human CD8^+^ T cells showed HLA-A*66:01-dependent, high-avidity recognition of HLA-A*66:01^+^ LCLs pulsed with these peptides (**Fig 5C**). We also used the COS-7 ORFeome transfection assay followed by candidate peptide selection with NetMHCpan to identify the KSHV epitope recognized by a fourth public TCR (TCR-24; **S1 Table**). This TCR was strongly associated with HLA-B*57 and was observed in 31 tumors from 5 PLWH and 4 HIV**^−^** individuals, all of whom expressed HLA-B*57:02 or HLA-B*57:03 (**S4B Fig**). Primary CD8^+^ T cells transduced with TCR-24 showed HLA-B*57:03-dependent, high-avidity recognition of the peptide corresponding to ORF59_171-179_ (**Fig 5D**).

### CD8^+^ T cells expressing KSHV-specific TCRs recognize HLA-matched PEL undergoing lytic replication of KSHV

To determine whether T cells expressing TCRs specific for KSHV lytic gene epitopes can recognize KSHV-infected cells presenting endogenously processed viral peptides, we tested a HLA-B*45:01^+^ primary effusion lymphoma (PEL) cell line carrying the KSHV genome, JSC-1, for expression of the ORF6_329-337_ epitope presented by HLA-B*45:01. Jurkat reporter T cells engineered to express either of our two ORF6_329-337_-specific TCRs showed low-level recognition of JSC-1 cells grown in the absence of activation signals (**Fig 6A**), which is likely accounted for by the observation that JSC-1 cells demonstrate lytic KSHV gene expression even in the absence of activation [49]. However, TCR-transduced Jurkat cells showed strongly enhanced TCR signaling when cultured with JSC-1 cells in which lytic replication had been chemically induced with 12-O-tetradecanoylphorbol-13-acetate (TPA), similar to the level of TCR signaling induced by direct CD3 stimulation with the CD3-specific monoclonal antibody OKT3. Primary CD8^+^ T cells engineered to express the ORF6_329-337_-specific TCRs also demonstrated IFN-γ production and cytotoxicity when cultured with TPA-treated JSC-1 cells (**Fig 6B, C**). Similar experiments with the HLA-B*45:01^+^ PEL cell line VG-1, which is profoundly deficient in TAP1 activity and antigen presentation [50], triggered weaker stimulation of Jurkat cells transduced with the ORF6_329-337-_specific TCRs.

**Figure 6.**
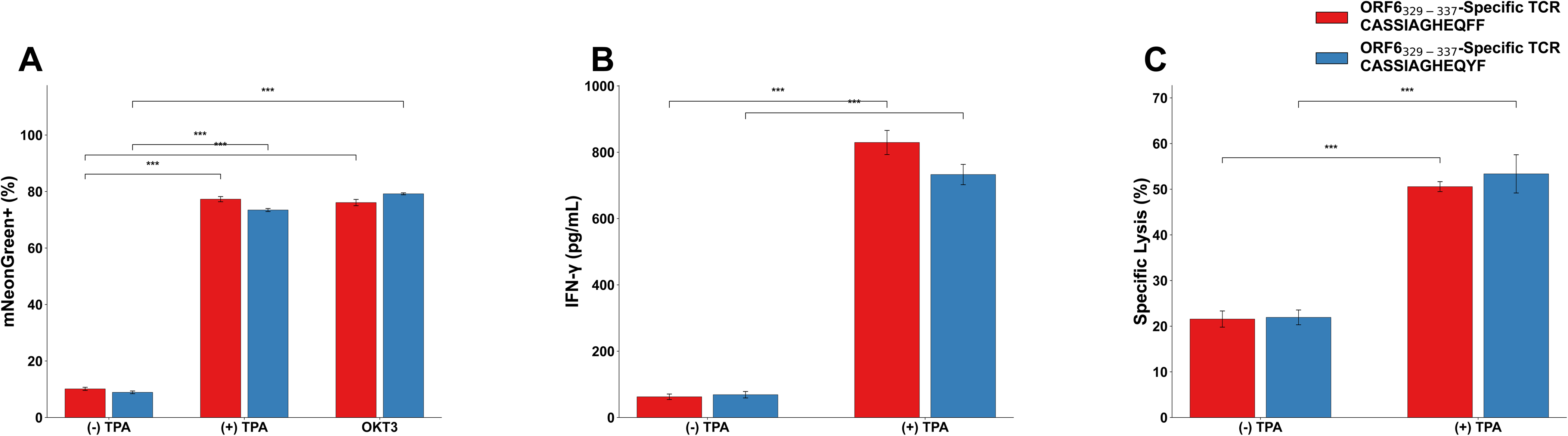
KSHV TCRs recognize endogenous KSHV peptides in KSHV^+^ PEL cells. **(A)** TCR signaling in Jurkat reporter cells (mNeonGreen fluorescence) co-cultured with ORF6_329-337_-specific TCR-2 (red) or TCR-3 (blue) in JSC-1 PEL cells with or without TPA treatment. OKT3 response, positive control. **(B)** IFN-γ production by primary human CD8^+^ cells transduced with ORF-6-specific TCRs cultured in the presence of JSC-1 PEL treated with or without TPA and **(C)** Cytotoxicity in a four-hour ^51^Cr release assay at an E:T ratio of 10:1. Mean +/- SEM. Two-sided t-test comparison of TPA-treated conditions to untreated controls, ***p ≤ 0.001.

### Lytic activation-dependent CD8^+^ T-cell recognition of KSHV-infected iTIME.219 endothelial cells

Finally, we examined whether T cells expressing KS TIL-derived KSHV-specific TCRs can recognize an endothelial cell line, iTIME.219, that carries a KSHV genome. The telomerase-immortalized iTIME.219 cell line carries a recombinant KSHV genome with a doxycycline-inducible KSHV transcription activator (RTA) gene that induces viral replication [51]. iTIME.219 cells constitutively express green fluorescent protein (GFP) when cultured in the absence of doxycycline and activation signals. Exposure of iTIME.219 cells to sodium butyrate and/or doxycycline, however, induces expression of RTA that in turn activates KSHV lytic gene expression as well as expression of red fluorescent protein (RFP), enabling both controlled induction and tracking of KSHV lytic reactivation.

We transduced iTIME.219 cells with lentiviral vectors encoding one of the three MHC class I alleles (HLA-B*45:01, HLA-A*66:01, HLA-B*57:03) that present the ORF6_329-337_, ORF57_381-389_, and ORF59_171-179_ peptides, respectively, to the KSHV-specific TCRs that we characterized. MHC class I-transduced, chemically induced iTIME.219 cells were co-cultured with TCR-transduced Jurkat reporter T cells to evaluate TCR signaling (**Fig 7A**). MHC-restricted TCR signaling was detected for all 4 TCR-peptide pairs tested, with TCR signaling only observed when TCR-transduced T cells were cultured with sodium butyrate- and doxycycline-treated iTIME.219 cells transduced with the appropriate MHC allele (**Fig 7B**). TCR signaling was also seen when TCR-transduced T cells were cultured with uninduced iTIME cells that had been transduced with the appropriate MHC allele and pulsed exogenously with the corresponding KSHV peptide. These findings confirm that T cells expressing TCRs specific for peptides encoded by KSHV lytic genes and presented by class I MHC alleles can recognize KSHV-infected endothelial cells presenting endogenously processed antigen.

**Figure 7.**
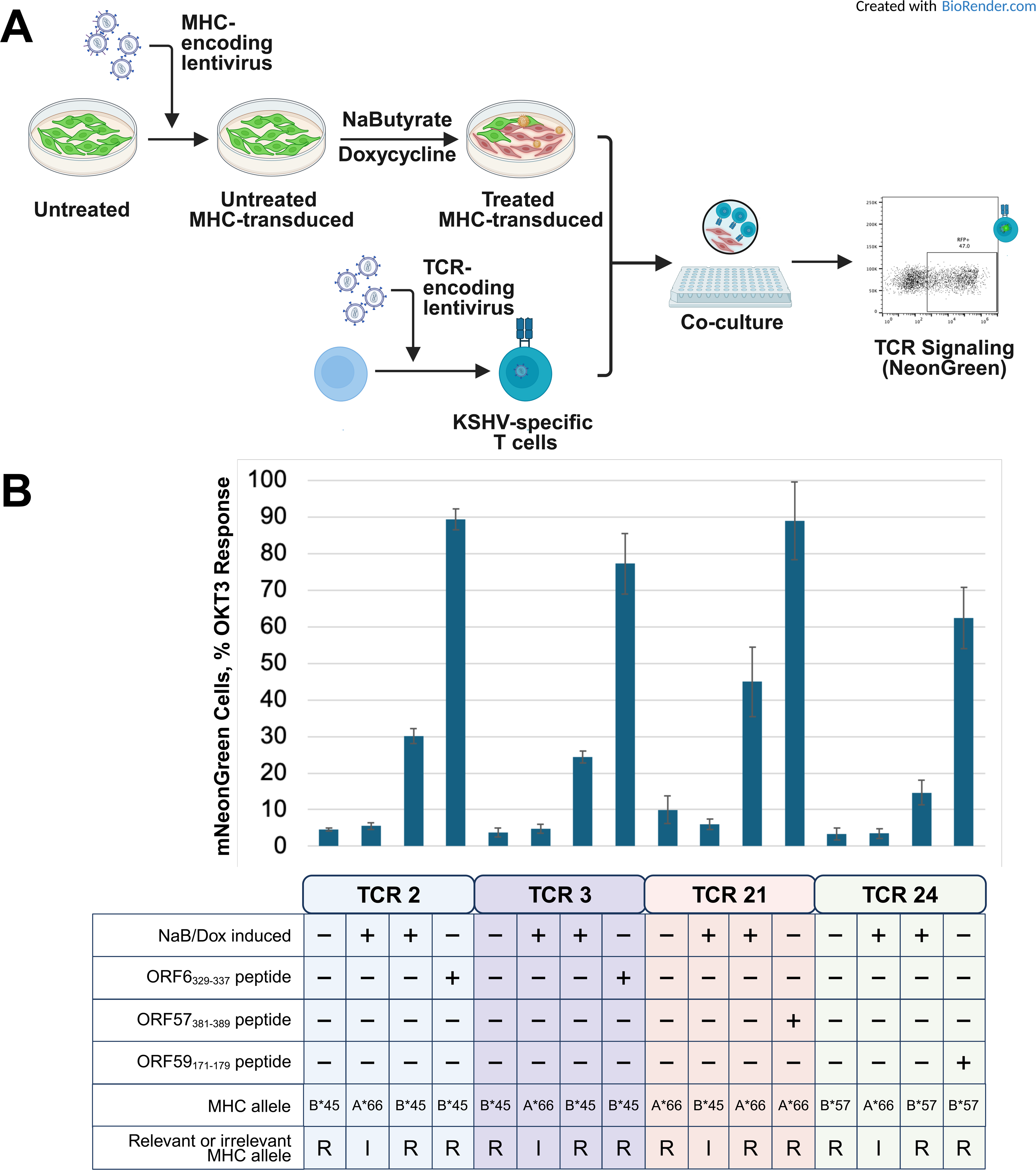
KSHV-specific TCRs recognize endogenous KSHV peptides on KSHV^+^ iTIME.219 cells. **(A)** Schematic representation of workflow used to evaluate recognition of KSHV-infected iTIME.219 endothelial cells by Jurkat reporter T cells transduced with KSHV-specific TCRs. **(B)** TCR signaling in Jurkat reporter cells (mNeonGreen fluorescence) transduced with the indicated TCR and co-cultured for 16 hours with iTIME.219 cells that were transduced with the indicated relevant or irrelevant class I MHC molecule and pre-treated with or with sodium butyrate / doxycycline (NaB/Dox) for 24 hours. Mean +/- pooled SD of triplicate determinations across at least two experiments.

## Discussion

In this study we identified T cells that commonly infiltrate KS tumors and carry αβ TCRs specific for peptides encoded by KSHV lytic genes and presented by class I MHC molecules. We further showed that the TCRs carried in these KS tumor-infiltrating lymphocytes specifically recognize KSHV-infected B-cells and endothelial cells presenting endogenously processed viral peptides. Of the 8 putative KSHV-specific TCRs we tested, 3 TCRs represent a shared, i.e., “public” T-cell response to KSHV, with two sharing specificity for ORF6_329-337_ presented by HLA-B*45:01, and one recognizing ORF59_171-179_ presented by HLA-B*57:03. The two ORF6-specific TCRs were detected in 29 distinct tumors from 14 HLA-B*45:01^+^ individuals in our KS cohort, while the ORF59-specific TCR was similarly detected in 31 distinct KS tumors from 10 HLA-B*57:03^+^ individuals. A fourth TCR was found to recognize ORF57_381-389_ presented by HLA-A*66:01 and HLA-A*66:02. This TCR was not observed in other individuals in our cohort but is notable because it comprised more than 23% and 43% of the TCRs in two non-contiguous tumors from an HLA-A*66:01^+^, HIV-seronegative participant with KS who subsequently achieved a complete response with therapy. One of the two public ORF6_329-337_-specific TCRs that we characterized was also observed in the same two non-contiguous tumors from this HIV-seronegative participant, who was also HLA-B*45:01**^+^**, and comprised 3.8% and 3.6% of the TCRs.

Our findings build on and extend previous investigations of the T-cell response to KSHV. Studies of asymptomatic KSHV-infected individuals and from individuals with KS have identified several peptides encoded by lytic or latent KSHV genes that can elicit *in vitro* cytokine responses from unsorted peripheral blood T cells or from purified CD8**^+^** or CD4**^+^** T cells [17–28, 30–32, 52–54]. Many of these previous studies focused primarily on peptides that are predicted to bind to HLA-A*02:01, which is the most common class I MHC allele carried in North American and European populations [55]. To our knowledge, our study is the first to identify and characterize specific αβ TCRs that recognize KSHV-encoded peptides with physiologically relevant low-nanomolar affinity. Most previous studies of the T-cell response to KSHV have utilized *in vitro* assays that cannot distinguish between high affinity/high specificity and low affinity/low specificity responses. Moreover, the TCRs we investigated were selected because they were observed in KS tumors from multiple individuals or were expressed on expanded TIL populations that massively infiltrated two KS tumors from an HIV-seronegative patient who achieved a complete response with therapy. The MHC restricting alleles associated with these TCRs are common in the general Ugandan population, which includes our cohort, but uncommon in North American and European populations [55]. Identification of the antigenic peptides recognized by other candidate KSHV-specific TCRs from our collection of >4,000 clustered unknown TCRs is actively underway.

Public TCRs are a common feature of herpesvirus-specific and other virus-specific T-cell repertoires [56–59]. The ORF6- and ORF59-specific TCRs characterized in this study are the first KSHV-specific public TCRs to be reported. Many herpesvirus-specific public TCRs recognize immunodominant viral antigens, such as CMV pp65_495-503_ [60] and EBV BMLF1_280-288_ [61]. Most are generated with high probability during T-cell receptor gene rearrangement, and their detection in many individuals is thought to reflect convergent recombination and selection. Although there is no evidence that herpesvirus-specific T cells bearing public TCRs have superior antiviral activity *per se*, their specificity is often associated with target proteins such as CMV pp65 and CMV IE-1 that are critical control points for viral replication [62]. The identification of public KSHV-specific TCRs such as those recognizing ORF6_329-337_ and ORF59_171-179_ in individuals with KS suggests that they may also be present in the KSHV-specific T-cell repertoire in asymptomatic KSHV-infected individuals, as well, and current studies in our lab are investigating this hypothesis.

The KSHV genes *ORF6*, *ORF57*, and *ORF59* are expressed during the lytic phase of the viral life cycle. The transcriptional profiling reported in this and our previous [33] study demonstrated that transcription of these and other lytic genes is consistently detected in KS tumors from both PLWH and HIV-seronegative individuals. Lytic reactivation is thought to modulate the KS tumor microenvironment and promote angiogenesis, inflammation, viral spread, and lesion progression [63, 64]. *ORF6*, *ORF57*, and *ORF59* represent the most abundant viral proteins expressed during viral reactivation in human endothelial cells [65]. *ORF6* encodes a single-stranded DNA binding protein that stabilizes viral DNA and enables efficient KSHV genome amplification, and *ORF59* encodes a viral DNA polymerase processivity factor that binds and stabilizes the viral DNA polymerase; *ORF6* and *ORF59* are both indispensable for viral DNA synthesis [66]. *ORF57* encodes a multifunctional RNA binding protein and post-transcriptional regulator that stabilizes viral RNA, regulates splicing, and promotes nuclear export and translation. Cytotoxic T cell targeting of KSHV-infected cells expressing *ORF6*, *ORF57*, and *ORF59* would be expected to inhibit viral replication by eliminating cells engaged in viral genome synthesis and the production of viral transcripts required for lytic infection.

Whether T cells recognizing the products of KSHV latent genes infiltrate KS tumors is a critically important question and an active focus of current investigation in our lab. The latent phase of the KSHV lifecycle is characterized by expression of a limited number of genes, including *LANA-1* (*ORF73*), *vCyclin* (*ORF72*), *vFLIP* (*ORF71*), *vIRF3* (*ORF10.5*), and *K12*. Our transcriptional profiling reveals high-level expression of *ORF71*, *ORF72*, and *K12* in KS tumors, and detection of LANA-1 protein by immunohistochemistry is a defining feature of KS tumor cells. LANA-1 contains a highly acidic, low complexity repeat region from residues ∼330 to ∼886 that inhibits both its efficient synthesis and its proteasomal degradation [67]. This region limits the entry of LANA-1 into the class I MHC antigen processing pathway and its subsequent presentation to CD8**^+^** T cells. CD4**^+^** T-cells with reactivity against peptides derived encoded by *LANA-1* and presented by class II MHC molecules have been described [21, 27, 28]. The EBNA-1 protein of EBV, to which LANA-1 is in many respects analogous, similarly inhibits its own proteasomal degradation and entry into the class I MHC antigen processing pathway through its Gly-Ala repeat region [68]. CD8**^+^** T-cell epitopes such as EBNA-1_407-417_ (HPVGEADYFEY) have been identified outside the Gly-Ala repeat region, however [69], and are recognized by a broad repertoire of TCRs including public TCRs [70, 71]. We hypothesize that LANA-1 also encodes antigenic peptides that are endogenously processed and presented by class I MHC molecules to CD8**^+^** T-cells, but anticipate that their identification will require innovative strategies. We are currently using such strategies to pursue the identification of CD8**^+^** T cell epitopes encoded by LANA-1 and other latent KSHV genes. Effective targeting of the large number of KS tumor cells that express only KSHV latent genes will be required to develop effective immunotherapeutic strategies for KS.

Our studies of the TIL in KS tumors from PLWH also identified several novel HIV-specific TCRs that showed high-avidity recognition of *Nef*- and *Vpr*-derived peptides. Two with shared TCRβ chain homology recognize Nef_71-79_ presented by HLA-B*42:01, with half-maximal lysis observed at peptide concentrations <1 nM, and one recognizing Vpr_34-42_ presented by HLA-B*42:01, with half-maximal lysis <5 nM. Alongside previously reported HIV Pol_982-990_ and Vpr_34-42_-specific TCRs [44], our newly identified HIV-specific TCRs were detected in epidemic KS tumors obtained both at baseline and from the same individuals after they had been on antiretroviral therapy (ART) for several months. These findings complement observations of HIV Nef protein in tumor-resident macrophages from Ugandan KS patients who failed to respond to ART despite undetectable plasma HIV viral loads [72], and suggest that HIV may persist in some refractory KS tumors where it continues to shape the tumor microenvironment and may contribute to poor treatment responses.

Comprehensive identification of T-cell responses to KSHV will lay the foundation for strategies to prevent and treat KS and other KSHV-associated diseases. These diseases most commonly develop in individuals with T-cell deficiency or dysfunction, such as in PLWH or recipients of allogeneic solid organ or hematopoietic cell transplants, typically many years after primary infection with KSHV [1–3, 14–16]. In these settings KS can remit following initiation of antiretroviral therapy or withdrawal of immune suppression. These observations suggest that loss or impairment of pre-existing KSHV-specific T-cell immunity contributes to the development of KSHV-related malignancies. Strategies that preserve or restore the T-cell component of KSHV-specific immunity in PLWH and others at risk should, therefore, have potential for the prevention or treatment of KSHV-associated disease. Once the major targets of the KSHV-specific T-cell response have been identified, multimer-based sorting strategies can be utilized to define the full repertoire of T cells recognizing virus-encoded epitopes. Profiling the transcriptomes, function, and antigen avidity of this repertoire in KSHV-seropositive children and adults, with or without KS, and with or without HIV, will elucidate the nature of the deficiencies that predispose to KS and other KSHV-associated diseases and will guide the development of actionable strategies to restore effective responses.

Cellular therapy with KSHV-specific T cells represents one such strategy that is already being investigated for the related oncogenic gammaherpesvirus EBV. Tabelecleucel is an off-the-<u>shelf</u> allogeneic EBV-specific T-cell therapy that is approved in the EU as standard of care for EBV-positive post-transplant lymphoproliferative disease [73, 74]. The ability to transfer specificity for KSHV-encoded antigens into healthy CD8**^+^** or CD4**^+^** T cells with lentiviral or non-viral vectors makes it theoretically possible to generate T-cell products with pre-specified composition that would be matched for the MHC alleles of the intended recipient and could target both lytic and latent KSHV antigens. TCR gene transfer could also aid in the design of such cellular therapy products by enabling preclinical studies with T cells from healthy donors to study the mechanisms by which KSHV-infected cells can escape immune recognition and elimination, and to identify how those mechanisms can be disabled or counteracted.

## Materials and Methods

### Subjects and samples

Adults 18 years or older with histologically confirmed Kaposi sarcoma who were naïve to antiretroviral therapy (ART) and had received no prior chemotherapy were enrolled between October 2012 and May 2021 on a prospective cohort study (termed “HIPPOS”) at the Uganda Cancer Institute (UCI) in Kampala, Uganda, as previously described [38]. Participants completed an enrollment visit and up to 10 follow-up visits over ∼1 year. At enrollment, participants completed a standardized medical history and physical examination, a blood sample and an oral swab were collected for KSHV testing, and peripheral blood mononuclear cells (PBMC) were isolated. HIV serology was performed, and CD4**^+^** T-cell and plasma HIV RNA testing was completed for HIV-seropositive participants. UCI medical charts were reviewed to record routine blood count data and chest X-ray findings. Full-thickness punch biopsies (4 x 4 mm diameter) of at least three non-contiguous cutaneous KS lesions and a punch biopsy of axillary skin not visibly involved with KS were also obtained and placed in either neutral buffered formalin or RNA*later* (Thermo Fisher Scientific, Waltham, MA, USA).

At follow-up visits, participants completed an interim medical history and detailed physical exam to assess treatment response and provided additional blood samples and oral swabs for KSHV testing. Up to three additional biopsies of non-contiguous KS lesions and a blood sample for PBMC isolation were obtained at follow-up visits at 3-, 36-, and 48-weeks following enrollment, and placed in RNA*later*.

Participants were provided chemotherapy at the UCI per standard guidelines independent of the study. First-line therapy consisted of combination bleomycin and vincristine (BV) or paclitaxel; both regimens were given every 3 weeks for six cycles. HIV-seropositive participants not already in HIV care were referred to local HIV clinics for management and initiation of ART according to Ugandan Ministry of Health guidelines.

### Clinical laboratory procedures

Plasma and oral swab samples were evaluated for KSHV DNA by quantitative, real-time PCR at the UCI-Fred Hutch Cancer Centre Laboratory in Kampala, Uganda as previously described [33, 38] Samples with >150 copies/mL of KSHV DNA were considered positive [75]. CD4**^+^** T-cell counts were measured by flow cytometry, and HIV-1 RNA levels were measured using real-time RT-PCR.

### RNA sequencing

Total RNA was extracted from cryopreserved KS tumor biopsies using the RNeasy Fibrous Tissue Mini Kit (QIAGEN, Venlo, The Netherlands), quantitated on a Qubit 3.0 Fluorometer (Thermo Fisher), and assessed on an Agilent 2200 Bioanalyzer (Agilent Technologies, Inc., Santa Clara, CA). RNA samples with RNA integrity number (RIN) greater than 7 were used for sequencing library preparation. Sequencing libraries from RNA extracted from cryopreserved KS tumor biopsies from 39 HIV-seropositive HIPPOS participants (epidemic KS) were prepared in the Fred Hutchinson Cancer Center (FHCC) Genomics Core Facility (Seattle, WA) using the TruSeq RNA Sample Prep Kit v2 (Illumina, San Diego, CA, USA), as previously described [33]. Library size distributions were validated using the Agilent 2200 TapeStation. Library QC, pooling of indexed libraries, and cluster optimization were performed using the Qubit 3.0 Fluorometer. Pooled libraries were sequenced on an Illumina HiSeq 2500 in “High Output” mode with a paired-end, 50-base read configuration. Sequencing libraries from RNA extracted from cryopreserved KS tumor biopsies from an additional 12 HIV-seronegative HIPPOS participants (endemic KS) were prepared and sequenced at Azenta (South Plainfield, NJ) using the NEBNext Ultra II RNA Library Prep Kit for Illumina (New England Biolabs, Ipswich, MA, USA). The sequencing libraries were validated on the Agilent TapeStation, quantified using the Qubit 3.0 Fluorometer as well as by quantitative PCR (KAPA Biosystems, Wilmington, MA, USA), and sequenced on a partial S4 lane of an Illumina NovaSeq 6000 with a paired-end, 150-base read configuration.

### Targeted gene expression analysis

RNA was extracted from formalin-fixed paraffin-embedded (FFPE) slides from 25 epidemic KS and 2 endemic KS tumor biopsies and 5 samples of uninvolved skin using the AllPrep DNA/RNA FFPE Kit (QIAGEN). Gene expression analysis was performed on RNA samples on the Nanostring platform (Nanostring Technologies, Seattle, WA, USA) using the Immunology V2 Panel supplemented with 30 user-defined viral and host gene probes (Supplementary Table S1). Of the 25 epidemic KS tumors profiled, 8 were also profiled using bulk RNA-Seq. Raw probe counts were normalized using the geometric mean of 15 housekeeping genes followed by analysis using nSolver software (Version 4.0) and the nSolver Advanced Analysis package (version 2.0.115).

### T-cell receptor β chain variable region sequencing

Genomic DNA was extracted from 363 cryopreserved biopsies of KS tumor and normal skin (NAT) using the DNeasy Blood & Tissue Kit (QIAGEN) and prepared for T-cell receptor β chain (*TRB*) variable region sequencing at survey level resolution [76] using the ImmunoSEQ hsTCRB v3.0 assay (Adaptive Biotechnologies, Seattle, WA). *TRB* sequencing was performed on pools of libraries from 11 - 15 tumor or skin samples on the Illumina MiSeq platform using a v3 150-cycle kit (Illumina, San Diego, CA) in the FHCC Genomics Core Facility (Seattle, WA) or the UCI-Fred Hutch Cancer Centre Laboratory (Kampala, Uganda).

### MHC genotyping

High-resolution genotyping of the MHC class I and class II loci of all participants in the epidemic and endemic KS cohorts and the Uganda BL cohort was performed on genomic DNA via next-generation sequencing as a contract service by Scisco Genetics, Seattle, WA. The full standardized protocol is available online [77]. Sequencing of the PCR amplicons spanning the MHC loci was performed on the Illumina MiSeq platform using a v2 500-cycle kit (Illumina, San Diego, CA) in the FHCC Genomics Core Facility or the UCI-Fred Hutch Cancer Centre Laboratory.

### Single-cell RNA sequencing (scRNA-Seq)

Viable cells were thawed from cryopreserved KS PBMC specimens and subjected to scRNA-Seq using the 10X Genomics (10X Genomics, Pleasanton, CA) platform. Single index chemistry was performed for all libraries using the following reagents to generate scRNA-Seq libraries for V1.0 chemistry: the Chromium^TM^ Single Cell 5’ Library Construction Kit, Chromium ^TM^ Single Cell A Chip Kit, ChromiumTM Single Cell 5’ Library & Gel Bead Kit, Chromium^TM^ Single Cell 5’ Feature Barcode Library Kit, and the Chromium^TM^ Single-Cell Enrichment, Human T-Cell kit.

For libraries generated using V1.1 chemistry, the Chromium^TM^ Next GEM Single Cell 5’ Library and Gel Bead Kit v1.1, and the Chromium^TM^ Next GEM Chip G Single Cell Kit were used.

Libraries were quantified using both the Qubit Fluorometer (Thermo Fisher, Waltham, MA), TapeStation using the Agilent TapeStation (Agilent, Santa Clara, CA) and by qPCR using the KAPA Library Quantification Kit for Illumina (Roche, Basel, Switzerland). Libraries were pooled and sequenced on the Illumina NovaSeq 6000 (Illumina, San Diego, CA) on an SP1 flow cell in the FHCC Genomics Core.

### RNA-seq analysis

A total of 5,355,792,915 reads were obtained across 110 bulk RNA-Seq libraries: 51 paired-end libraries (4,049,586,334 reads; mean 79,403,654 reads per sample) generated in this study from 39 epidemic KS and 12 endemic KS tumor biopsies (cohort “Hippos”), and 59 single-end libraries (1,306,206,581 reads; mean 22,139,095 reads per sample) re-analyzed from Tso *et al*. ([34]; n = 8) and Lidenge *et al*. ([35]; n = 51). Raw reads from the published validation cohorts were retrieved from SRA using the NCBI SRA-Toolkit. All libraries were processed through a single nf-core/rnaseq pipeline run (v3.19.0; [78]) using the Docker execution profile. Adapter contamination and low-quality bases were removed with Trim Galore v0.6.10 [79], retaining the pipeline’s default parameters (Phred quality cutoff Q20, minimum read length 20 bp, automatic adapter detection). A combined human + viral reference was constructed by concatenating the GENCODE v47 primary-assembly human reference (GRCh38.primary_assembly.genome.fa; gencode.v47.primary_assembly.annotation.gtf) with the NCBI RefSeq genomes for KSHV/HHV-8 (GCF_000838265.1) and HIV-1 (GCF_000864765.1). The viral GFF3 annotations were converted to GENCODE-compatible GTF and gene/transcript identifiers were prefixed with KSHV_ and HIV1_ to prevent ID collisions with the human annotation. The combined annotation is deposited as (Supplementary File S1; rnaseq/GRCh38_KSHV_HIV1.gtf.gz). This GTF, together with the GRCh38 + KSHV + HIV-1 concatenated FASTA, was used to build a STAR v2.7.11b genome index, and trimmed reads were aligned with STAR in two-pass mode using the pipeline’s default ENCODE-style parameters. Transcript-level abundance was estimated from STAR’s transcriptome-coordinate BAMs with Salmon v1.10.3 [80] in alignment-based mode (library type IU), using the deposited combined GTF as the gene–transcript map. Salmon quant.sf and quant.genes.sf outputs were aggregated into per-sample transcripts-per-million (TPM) matrices spanning ∼78,932 human, 96 KSHV and 10 HIV-1 genes, with samples as rows, gene IDs as columns and TPM values in each cell. KSHV gene IDs were mapped to common gene names and lytic-cycle stage using a curated lookup derived from the HHV-8 RefSeq GFF (Supplementary File S1; see Zenodo file rnaseq/kshv_gene_map.tsv).

Quality metrics (FastQC v0.12.1, Picard MarkDuplicates v3.1.1, Qualimap RNA-Seq v2.3, RSeQC v5.0.2, dupRadar v1.32.0, samtools v1.21) were aggregated with MultiQC, and downstream count-level QC was performed with DESeq2 v1.28.0 within the same pipeline. We further included gene-level TPM matrices for 30 sun-exposed (lower-leg) and 30 non-sun-exposed (suprapubic) skin samples from GTEx v8 (https://gtexportal.org/home/) as a baseline transcriptomic profile of normal skin. The harmonized TPM matrices were then formatted for analysis with CIBERSORTx [36, 37], and the immune cell-type composition of each sample was estimated against the LM-22 leukocyte signature via the CIBERSORTx web interface (https://cibersortx.stanford.edu/).

### T-cell receptor repertoire analysis

T-cell repertoire analyses were conducted using the R statistical programming language (Team, 2021). A reproducible workflow of all steps in data QC, data transformation, and analysis was developed using Jupyter notebooks and the Nextflow workflow management system [81]. Repertoire analysis was performed with open-source R packages and custom scripts were implemented in LymphoSeq2, an R package that we developed for the exploration of Adaptive Immune Receptor Repertoire Sequencing (AIRR-Seq) data [82].

To investigate intra-tumoral T-cell heterogeneity, we sequenced a total of 474 TCR repertoires from both tumor and adjacent normal tissues collected from 144 individuals enrolled in the study. Repertoires with less than 1,000 total *TRB* sequences were removed from the analysis, leaving a total of 363 TCR repertoires from 299 KS tumor biopsies (221 epidemic KS, 78 endemic KS) and 64 samples of uninvolved axillary skin (39 epidemic KS, 25 endemic KS), and 7 samples from single-cell tumor suspensions subjected to a Rapid Expansion Protocol (6 epidemic KS and 1 endemic KS).

### Prediction of T-cell antigenic specificity and MHC restriction

The TRB CDR3 amino acid sequences in each repertoire were compared with the VDJdb [40–42] and McPAS-TCR [43] databases of annotated TRB sequences, with high confidence of antigenic specificity (VDJdb score >1; McPAS-TCR antigen identification method < 3), to identify sequences that have previously been associated with a T-cell response to a specific microbial or tissue antigen or a specific pathological condition. TCRs from public databases were filtered to ensure that the recorded MHC allele in the database was also found in the individual with the sequence of interest.

Antigenic specificity groups of T-cell receptors carrying specific TRB CDR3 amino acid sequences were computationally predicted using GLIPH2 [45]. Sequences from T-cell receptor repertoires were labeled with repertoire ID and phenotype (epidemic KS, endemic KS). MHC typing information from the KS cohort was formatted as per GLIPH2 requirements. A reference database for GLIPH2 was generated using 35 control AIRR-Seq PBMC datasets from sub-Saharan Africa. The custom reference catalogued the CDR3 length distribution, V-gene usage, and CDR3 composition of 932,116 unique TRB CDR3 sequences.

TRB CDR3 amino acid sequences with likely antigenic specificity for CMV, EBV, HIV, *Mycobacterium tuberculosis* (*Mtb*), or an identified tissue antigen, were identified by first finding all specificity groups that contained an annotated TRB sequence reported in one or more of the public databases. All TRB CDR3 amino acid sequences belonging to these specificity groups were identified as sequences with likely antigenic specificity for CMV, EBV, HIV, *Mtb*, a specific tissue antigen, or pathological condition. GLIPH2 groups that did not contain any TRB CDR3 amino acid sequences with associated antigenic specificity were categorized as “Clustered Unknowns”. Sequences not grouped into antigenic specificity groups by GLIPH2 and with yet undetermined antigenic specificity were categorized as “Unclustered Unknowns.”

### Analysis of single-cell RNA-seq gene expression (GEX) and VDJ libraries

We built a structured analysis pipeline using NextFlow to process scRNA-Seq GEX + VDJ sequencing libraries from PBMC samples of 20 individuals with epidemic and endemic KS. We quantified the number of cells across both GEX and VDJ libraries using the Cell Ranger multi (pipeline version: Cell Ranger-7.0.1). The references for Cell Ranger GEX (Human reference (GRCh38) - 2020-A) and VDJ (Human V(D)J reference (GRCh38) ensembl-7.0.0) were obtained from the 10X Cell Ranger website (https://www.10xgenomics.com/support/software/cell-ranger/downloads). For quality control and preprocessing, we utilized the Bioconductor package scuttle to compute per-cell metrics [83]. We used the isOutlier function from the Bioconductor package scater to identify outliers based on per-cell metrics, with a threshold of 5 median average deviations [83]. Cell counts were normalized, and clustering was performed using scuttle and scater [83]. Doublet clusters were identified based on gene expression using the scDblFinder [84]. Seuratv4 [85], and scRepertoire [86] were employed to merge gene expression and VDJ data for each sample. T-cell sequence data from VDJ libraries were incorporated into the metadata field of the Seurat objects.

Gene expression and VDJ data from the 20 PBMC single-cell datasets were integrated using the Seurat rpca protocol [https://satijalab.org/seurat/articles/integration_rpca.html]. Ensembl IDs were replaced with gene symbols to prepare the integrated object for cell type classification.

### Antigen grouping of single cell RNA-seq VDJ libraries

GLIPH2 results from the bulk TRB sequencing data were merged with TRB sequences from the single-cell datasets to assign cells to antigenic specificity groups. This merged table was then filtered to retain only those TRB sequences classified as “Clustered Unknowns” (as defined in the previous section). For each retained TRB sequence, the paired TRA sequence was extracted from the single-cell repertoire data.

### TCR gene synthesis, lentiviral production, and T-cell transduction

Paired TRA and TRB contig sequences obtained from 10x Genomics scRNA-seq VDJ libraries were used alongside reference V, J, and C gene segment sequences from the International Immunogenetics Information System (IMGT) Repertoire to assemble full-length TCR constructs for gene synthesis [87–90]. Bulk TRA and TRB AIRR-seq data were used to infer hyperexpanded clonotypes, with IMGT repertoire locus gene sequences used to construct full-length TCR gene synthesis products. The amino acid sequences of the TCR α and β chains carried by the 9 KSHV-or HIV-specific TCRs that were studied, along with their inferred peptide specificity and class I MHC restriction, are detailed in Supplementary Table S2. Codon-optimized, full-length αβ TCRs were designed to contain a Kozak sequence, followed by the TCRβ chain, a self-cleaving porcine teschovirus-1 2A sequence, and the TCRα chain, with cysteine point mutations in the constant regions to crosslink each αβ chain and minimize mispairing with endogenous TCRs. Lentiviral transfer plasmids encoding each αβ TCR were synthesized in the pRRLSIN.cPPT.MSCV/WPRE backbone under control of the murine stem cell virus (MSCV) promoter (Azenta Life Sciences), and recombinant lentiviruses were produced by the FHCC Vector Production Facility.

Lentiviral supernatant was titered in NR4A1-mNeonGreen Jurkat clone E6.1 cells [46]. Transduction efficiency was determined as the percentage of DAPI^−^ TCR^+^ cells by flow cytometry using a PE-conjugated anti-human αβ TCR antibody (Miltenyi Biotec) at ≥3 days post-transduction. For functional KSHV-reactivity screens using Jurkat reporter cells, 1-2 µL lentivirus per 5 × 10^5^ cells in CTL medium (RPMI-1640 containing HEPES and L-glutamine (Thermo Fisher Scientific) supplemented with 10% heat-inactivated human serum (Bloodworks Northwest), an additional 2 mM L-glutamine, 1% penicillin/streptomycin (Thermo Fisher Scientific) and 50 μM β-mercaptoethanol (Millipore Sigma) containing 4.5 µg/mL polybrene (Millipore Sigma) was added and cells were spinoculated at 800 x g for 90 minutes at 32 ⁰C in a 48-well plate format.

Cells were washed and resuspended in fresh CTL media one day post-transduction and assessed for TCR surface expression by flow cytometry at a minimum of 3 days post-transduction. For each TCR, maximal signaling capacity was established by 24-hour stimulation with 30 ng/mL anti-CD3 (OKT3, Invitrogen), with signaling quantified by flow cytometric mNeonGreen emission.

Human CD8^+^ T cells were isolated from healthy donor PBMCs using the Miltenyi Biotec CD8^+^ T Cell Isolation Kit. For primary CD8^+^ T-cell transductions, 5 × 10^5^ cells in a 48-well plate were activated with CD3/CD28 Dynabeads (Thermo Fisher Scientific) at a 3:1 ratio overnight in 1 mL CTL medium supplemented with 50 U/mL IL-2 (PeproTech). Lentivirus was added (1-2 µL per 5 × 10^5^ cells), and cells were spinoculated at 800 x g for 90 minutes at 32 ⁰C, followed by a half-media change after 8 hours. Dynabeads were removed 3 days post-activation, and cells were expanded to a 24-well plate with 2.45 mL CTL medium containing IL-2 containing. On Day 5, cells were split into two wells of a 12-well plate with 2.45 mL CTL medium containing IL-2. On Day 6, 1 mL fresh CTL + IL-2 was added, and cells were maintained with half-media changes every other day until use ≥9 days post-transduction. Transduction efficiency was confirmed using tetramer staining with fluorochrome-conjugated tetramers (e.g., APC-labeled Vpr_34-42_-tetramer) when available.

### Class I MHC plasmid synthesis

Class I MHC mammalian expression vectors were generated either by *de novo* gene synthesis (Azenta Life Sciences) or obtained from the Addgene plasmid repository. For synthesized constructs (HLA-A*02:01, HLA-B*45:01), IMGT reference HLA allele sequences were assembled with a Kozak sequence at the 5’ end and native 3’ stop codon. Codon-optimized sequences were cloned into the pcDNA3.1(+) expression vector (Thermo Fisher Scientific). Addgene-derived plasmids (HLA-A*66:01, #203291; HLA-B*57:03, #203260) were a gift from

Derin Keskin and Catherine Wu [91]. All plasmids were sequence-verified before use in downstream transfection assays.

### KSHV plasmids and minigenes

A complementary DNA (cDNA) library of all annotated protein-coding ORFs encoded by the KSHV genome (92 total constructs) was generated in the pDEST103 plasmid backbone [92]. A modified LANA construct (ΔLANA) containing the first 1,035 and final 735 nucleotides was synthesized, omitting the long acidic repeat region and insertion of a Kozak consensus sequence at the 5’ terminus (Azenta Life Sciences). To generate sufficient DNA for downstream transfection assays, the resulting 93 constructs were transformed into DH5α bacterial cells and expanded to scale. Plasmid DNA was isolated and quantified using a Qubit 3.0 Fluorometer.

KSHV ORF minigene expression constructs were generated by PCR amplification of segments from the full-length ORF plasmid, followed by directional cloning. HLA-binding prediction tools (NetMHCpan-4.0 [48], SYFPEITHI [93]) were used to guide primer design for minigene construction, ensuring that strong binders for the respective HLA alleles were not truncated. Minimal epitope expression plasmids for ORF6 and ORF57 were synthesized *de novo* as PAGE-purified duplex oligonucleotides (IDT) designed to include a 5’ Kozak sequence, in-frame stop codon, and 3’ overhanging deoxyadenosines to facilitate TOPO TA cloning. Constructs were cloned into the pcDNA3.1/V5-His TOPO TA mammalian expression vector (Invitrogen, Thermo Fisher Scientific). Colony PCR was performed on 4-6 bacterial transformants per construct, and clones producing amplicons of the expected band size were confirmed by Sanger sequencing. Plasmid DNA concentrations were measured on a Qubit 3.0 Fluorometer prior to transfection assays.

### Transient transfections in COS-7 cells

For initial identification of TCR-stimulatory ORF-MHC conditions, COS-7 cells were co-transfected with either (i) pools of plasmids encoding 9-12 KSHV ORFs with a plasmid encoding the predicted MHC class I restricting alleles for each TCR, or (ii) ORF plasmid pools alone. Iterative screening was performed with individual ORFs or minigene constructs in the presence or absence of the predicted MHC class I restricting allele. Transfections were performed in 96-well flat-bottom plates seeded the previous day with 9 × 10^3^ COS-7 cells in antibiotic-free medium. Each well received 150 ng of plasmid DNA and FuGENE6 transfection reagent (Promega) at a 3:1 reagent: DNA mass ratio. At 24 hours, HLA-A*02:01 expression (control) was confirmed by flow cytometry using a PE-conjugated anti-human HLA-A2 antibody (BD Pharmingen). Cells were then resuspended in CTL medium in preparation for co-culture with TCR-transduced Jurkat reporter T cells.

### TCR signaling assays

For Jurkat co-cultures with transfected COS-7 cells, 9 × 10^3^ TCR transduced cells were added to each well of a 96-well plate containing COS-7 cells in CTL medium. Peptides were synthesized by GenScript, dissolved in DMSO at a stock concentration of 10 mg/mL, and added to target cells at the indicated final concentrations (nM) before T-cell addition. After 16 hours, Jurkat reporter T cells were collected from each well, stained with DAPI (1 µg/mL), and assessed for mNeonGreen expression by flow cytometry. HLA-A*66:01, HLA-B*42:01, HLA-B*45:01, and HLA-B*57:03 LCL cells established by immortalization with EBV-supernatant and 50 µg/mL cyclosporin A (Millipore Sigma) as previously described [94]. Co-cultures were performed in 96-well flat-bottom plates in triplicate, with 5 × 10^3^ target cells per well and a 10:1 effector:target (E:T) ratio. The number of Jurkat reporter T cells plated was adjusted based on transduction efficiency. For peptide-containing conditions, peptides were added as above. For maximal signaling controls, OKT3 (30 ng/mL, Thermo Fisher Scientific) was added to Jurkat only wells. After 16 hours, co-cultures were harvested, stained with DAPI (1 µg/mL), and assessed for mNeonGreen expression by flow cytometry.

### Cytotoxicity assays

1 × 10^6^ target cells (LCL or PEL) were labeled overnight at 37°C with 100 µCi of ^51^Cr, washed twice, resuspended in CTL medium, and plated in triplicates (5 × 10^5^ cells/well) in 96-well round-bottom plates. For peptide conditions, peptides were serially diluted from 10 mg/mL stock concentrations into CTL medium to achieve the indicated final concentrations (nM). Primary CD8^+^T cells transduced to express each HIV- or KSHV-specific TCR were added at a 10:1 E:T ratio. Detergent was used as a positive control (maximum lysis) and media as a negative control (spontaneous lysis). After 4 hours, 30 µL of supernatant was harvested from each well, transferred to a 96-well LumaPlate, dried overnight, sealed, and read on a MicroBeta microplate counter (PerkinElmer) using TopCount software.

### IFN-γ enzyme-linked immunosorbent assays (ELISAs)

Supernatants from 16-hour co-cultures of 5 × 10^5^ target cells (± peptide) with TCR transduced primary CD8^+^ T cells at a 10:1 E:T, plated in triplicate, were assayed for IFN-γ by ELISA, according to the manufacturer’s protocol (BD Biosciences). Absorbance was measured on a microplate reader using Gen5 v1.11 software. Standard curves were generated by linear regression analysis of serial dilutions of recombinant IFN-γ, and the resulting equation was used to calculate IFN-γ concentration in experimental co-culture conditions.

### PEL cell culture and induction of lytic replication

VG-1 and JSC-1 were generously gifted by Dr. Denise Whitby at the Frederick National Laboratory for Cancer Research. An additional aliquot of VG-1 was generously provided by Dr. Eva Gottwein at the Department of Microbiology-Immunology, Northwestern University. To chemically induce lytic KSHV replication, PEL cells were treated with 12-O-tetradecanoylphorbol-13-acetate (TPA; Cell Signaling Technology), as previously described [49]. Briefly, cells were resuspended at a density of 3 × 10^5^ cells/mL in RPMI-1640 medium supplemented with 20% FBS, containing 20 ng/mL TPA, prepared from a 200 µM working stock in DMSO. Cells were incubated for 24h, then collected and transferred into fresh medium before flow cytometry staining or functional co-culture assays.

### KSHV-infected iTIME.219 endothelial cell lentiviral transduction, lytic induction, and co-culture

An inducible telomerase-immortalized endothelial cell line latently infected with recombinant KSHV.219 (iTIME.219), was obtained from BEI Resources (NR-56694) [46] iTIME.219 cells were cultured in Vasculife Basal Medium supplemented with cytokines in the Vasculife SMV Lifefactor kit (Fisher Scientific). Lentiviral transfer plasmids encoding HLA-A*66:01 or HLA-B*45:01 were synthesized in the pRRLSIN.cPPT.MSCV/WPRE backbone under control of the murine stem cell virus (MSCV) promoter (Azenta Life Sciences), and recombinant lentiviruses were produced by the FHCC Vector Production Facility. Lentiviral supernatants were titered in HLA-low NR4A1-mNeonGreen Jurkat clone E6.1 cells, and transduction efficiency was determined as the percentage of DAPI^−^, MHC-I^+^ cells by flow cytometry using a PE-conjugated anti-human HLA-ABC antibody (W6/32, BD Pharmingen) at ≥3 days post-transduction. The iTIME.219 cells were transduced with 2 µL lentivirus per 5 × 10^5^ cells in medium containing 4.5 µg/mL polybrene (MilliporeSigma), and spinoculated at 800 x g for 90 minutes at 32 ⁰C in a 48-well plate format. A full media change was performed one day post-transduction.

To induce lytic replication, iTIME.219 cells were seeded at 3.5 × 10^5^ cells per well in a 6-well plate containing 2 mL/well complete growth medium and the following day were treated with 1 µg/mL doxycycline and 1 mM sodium butyrate (Thermo Fisher Scientific) Lytic induction was confirmed 24hrs post-treatment by flow cytometric detection of RFP expression, with untreated, latently infected cells used as negative controls.

### Flow cytometry

Cells were stained on ice with fluorochrome-conjugated monoclonal antibodies against αβ TCR (PE; Miltenyi Biotec). After staining, cells were washed and counterstained with DAPI (1 µg/mL) for live-dead discrimination. Flow cytometry was performed at the FHCC Flow Cytometry Shared Resource. Cells were gated sequentially on FSC-A versus SSC-A, DAPI-negative events, FSC-A versus FSC-H to exclude doublets, and fluorochrome-positive events (*e.g.*, mNeonGreen^+^). Data were analyzed using FlowJo software.

### Statistical analysis

GraphPad Prism v10 was used for statistical analysis and to generate TCR signaling, CRA, and ELISA figures. P values were calculated using a two-sided Student’s t-test.

## Data Availability

AIRR-Seq datasets from the 363 KS tumor and NAT samples are publicly available for access on the Adaptive Biotechnologies immunoSEQ Analyzer portal at (https://clients.adaptivebiotech.com/pub/ravishankar-2024-jem). All R scripts, metadata, GLIPH2 references and Nextflow workflows to analyze and visualize the data used in the study are available on GitHub (https://github.com/shashidhar22/ks_manuscript). Data generated by Nextflow pipelines and GLIPH2 used to generate figures for the manuscript are available on Zenodo (https://zenodo.org/records/10594758). RNA-Seq datasets from 51 KS tumors and scRNA-seq GEX+VDJ datasets from 20 KS PBMC libraries are available through the NCBI under BioProject number PRJNA1068629 (https://www.ncbi.nlm.nih.gov/bioproject/PRJNA106 8629).

## Ethics Statement

The studies involving human participants were reviewed and approved by the Makerere University School of Medicine Research and Ethics Committee, the Fred Hutchinson Cancer Center IRB, and the Uganda National Council for Science and Technology. All participants provided documentation of informed consent.

## Author Contributions

Conceptualization: SR, AMHT, ILT, DMK, WTP, EHW

Data curation: PM, JN, DM, SS, WTP, KRM, AMHT, ILT, CPM, WTP, EHW

Formal analysis: SR, ILT, CPM, WTP, EHW

Funding acquisition: WTP, EHW

Investigation: SR, AMHT, ILT, CPM, PM, JN, JK, DM, SS, LDA, DGC, LO, PA, JO, DMK, LJ, WTP, EHW

Methodology: SR, AMHT, ILT, CPM, DMK, LJ, WTP, EHW

Project administration: SR, AMHT, ILT, PM, JO, WTP, EHW

Resources: JW

Software: SR

Supervision: DMK, WTP, EHW

Validation: SR, AMHT, ILT, CPM, WTP, EHW

Visualization: SR, CPM, EHW

Writing – original draft: SR, EHW

Writing – review & editing: SR, AMHT, ILT, CPM, DMK, WTP, EHW

## Acknowledgments

We would like to acknowledge the HIPPOS study team, the Data and Laboratory Teams at the Hutchinson Center Research Institute – Uganda, HIPPOS study participants, and their families for enabling this research. We thank Dr. Jürgen Haas and Dr. Michael Stürzl for providing KSHV ORFs for cloning into the KSHV ORF library, Dr. Jeffrey Vieira, Sydney Favors, Denise Tong, and Dr. Kerry J. Laing for assistance with generation and maintenance of the KSHV ORF library, Carlissa J. Burrow for assistance with the COS-7 transfection assay, Dr. Phillip D. Greenberg for providing the TCR lentiviral vector backbone (pRRLSIN.cPPT.MSCV/WPRE), Dr. Aude G. Chapuis for providing the Jurkat reporter T cells, Dr. Denise Whitby and Dr. Eva Gottwein for providing JSC-1 and VG-1 cells, and Graden Chan and Naomi Yoon for technical assistance with the iTIME.219 experiments. Finally, we would like to thank The Genotype-Tissue Expression (GTEx) Project for data used in this manuscript: the Sun-exposed skin and Non-sun-exposed skin datasets from the v8 data release.

## Supporting information

**S1 Table.**
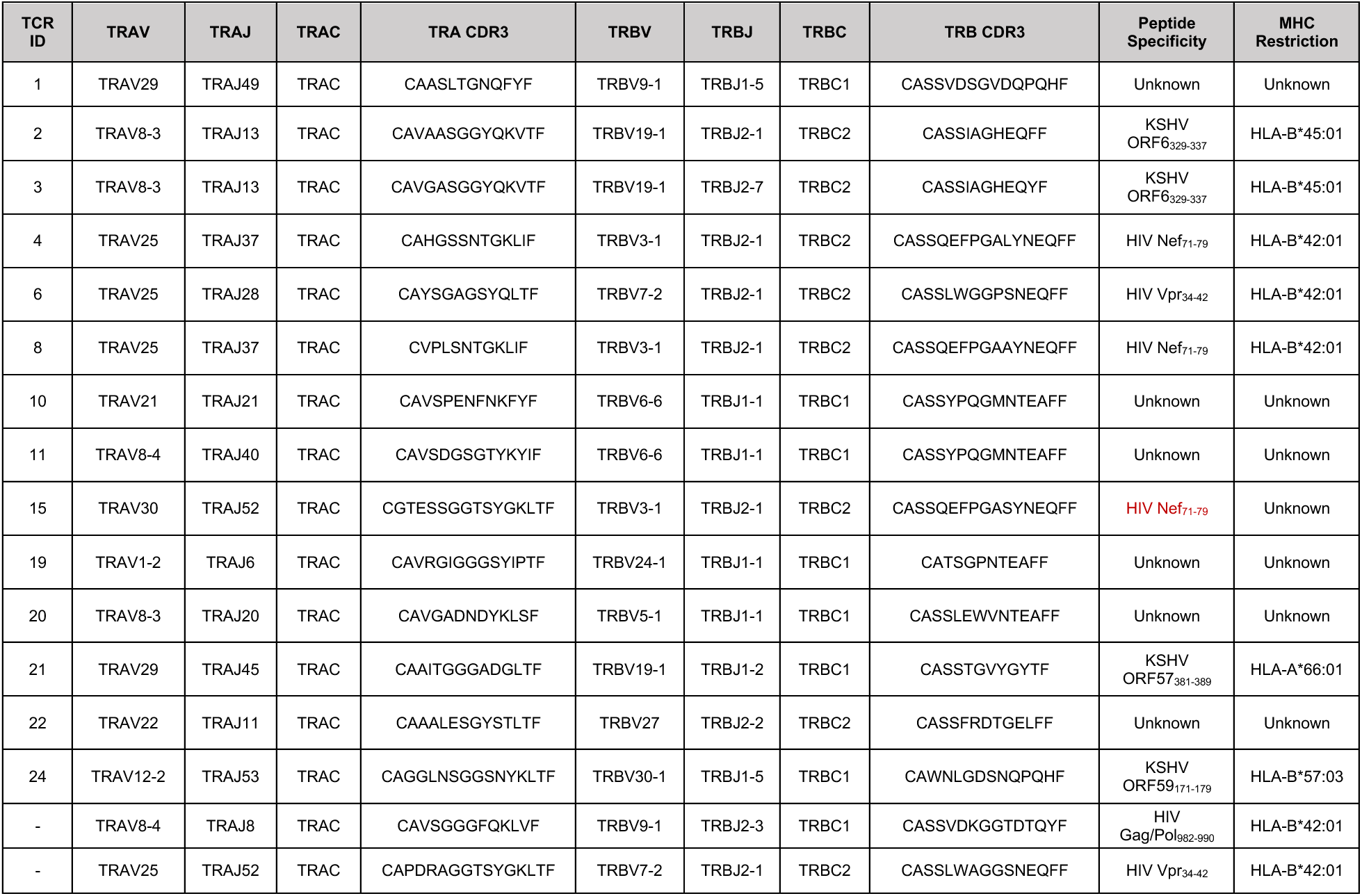
T-cell receptors investigated in this study.

**S2 Table.** HIV-encoded peptides known to be recognized by HLA-B*42:01-restricted CD8^+^ T cells [47].

| Peptide | Source protein/residues |
| --- | --- |
| FPRPWLHGL | Vpr<sub>34-42</sub> |
| HPKVSSEVHI | Vif<sub>48-57</sub> |
| IIKDYGKQM | Pol<sub>982-990</sub>; Integrase<sub>267-275</sub> |
| IPRRIRQGF | Env<sub>843-851</sub>; gp41<sub>332-340</sub> |
| LPPIVAKEI | Pol<sub>743-751</sub>; Integrase<sub>28-36</sub> |
| RLRPGGKKHY | Gag<sub>20-29</sub>; p17<sub>20-29</sub> |
| RPNNNTRKSI | Env<sub>298-307</sub>; gp120<sub>298-307</sub> |
| RPQVPLRPM | Nef<sub>71-79</sub> |
| TPGPGVRYPL | Nef<sub>128-137</sub> |
| TPQDLNTML | Gag<sub>180-188</sub>; p24<sub>48-56</sub> |
| YPGIKVKQL | Pol<sub>426-434</sub>; RT<sub>271-279</sub> |

**S1 Fig.**
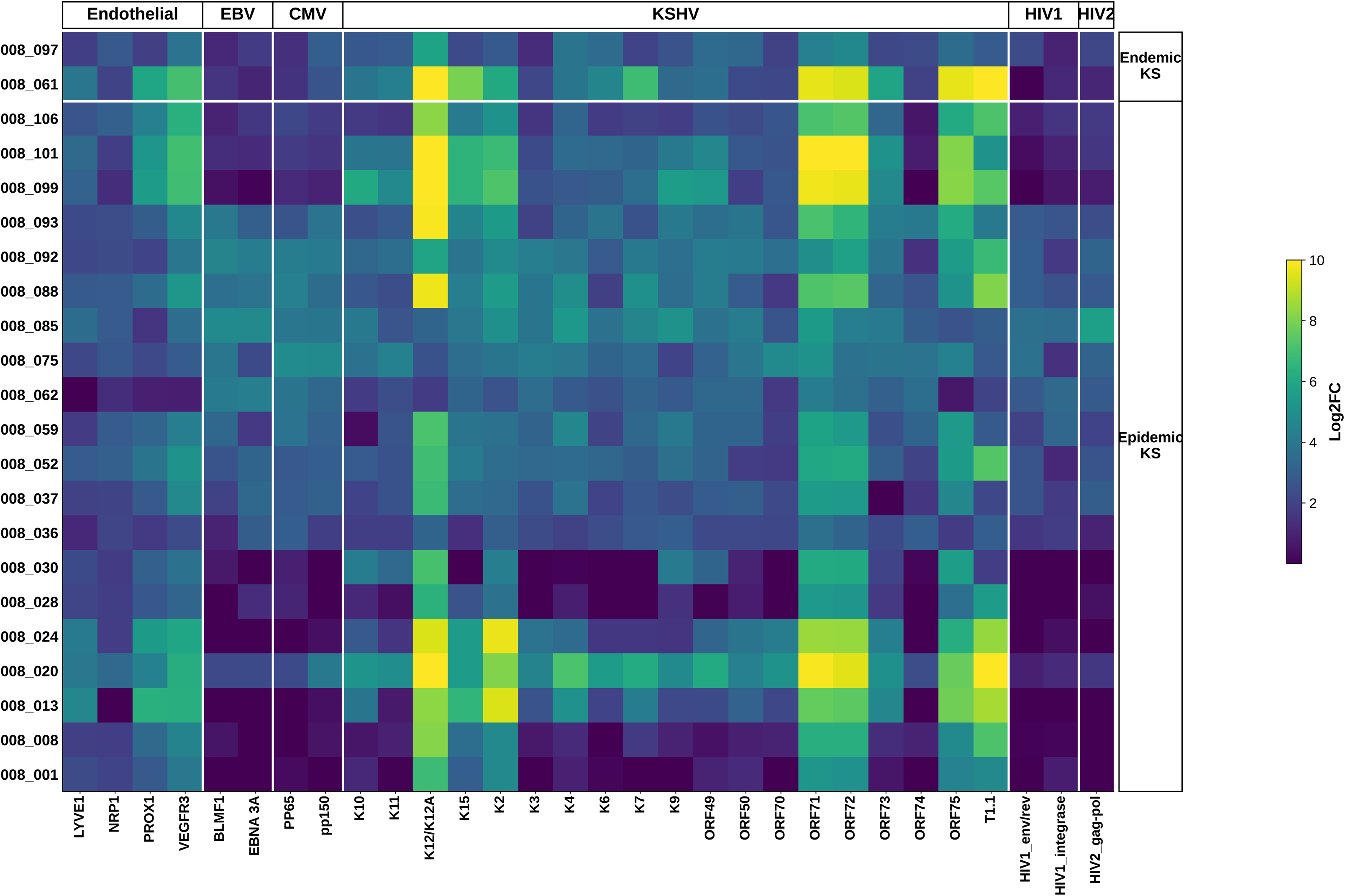
Targeted gene expression profiling of epidemic and endemic KS tumors. (**A)** Heatmap of expression of select KSHV genes in 2 tumors from individuals with endemic KS (top) and 20 tumors from individuals with epidemic KS (bottom) determined with a targeted gene expression platform. The log2 fold-change between average expression of each gene in the KS tumors and in 5 samples of clinically uninvolved skin from the axilla (NAT) is represented in color according to the legend at far right.

**S2 Fig.**
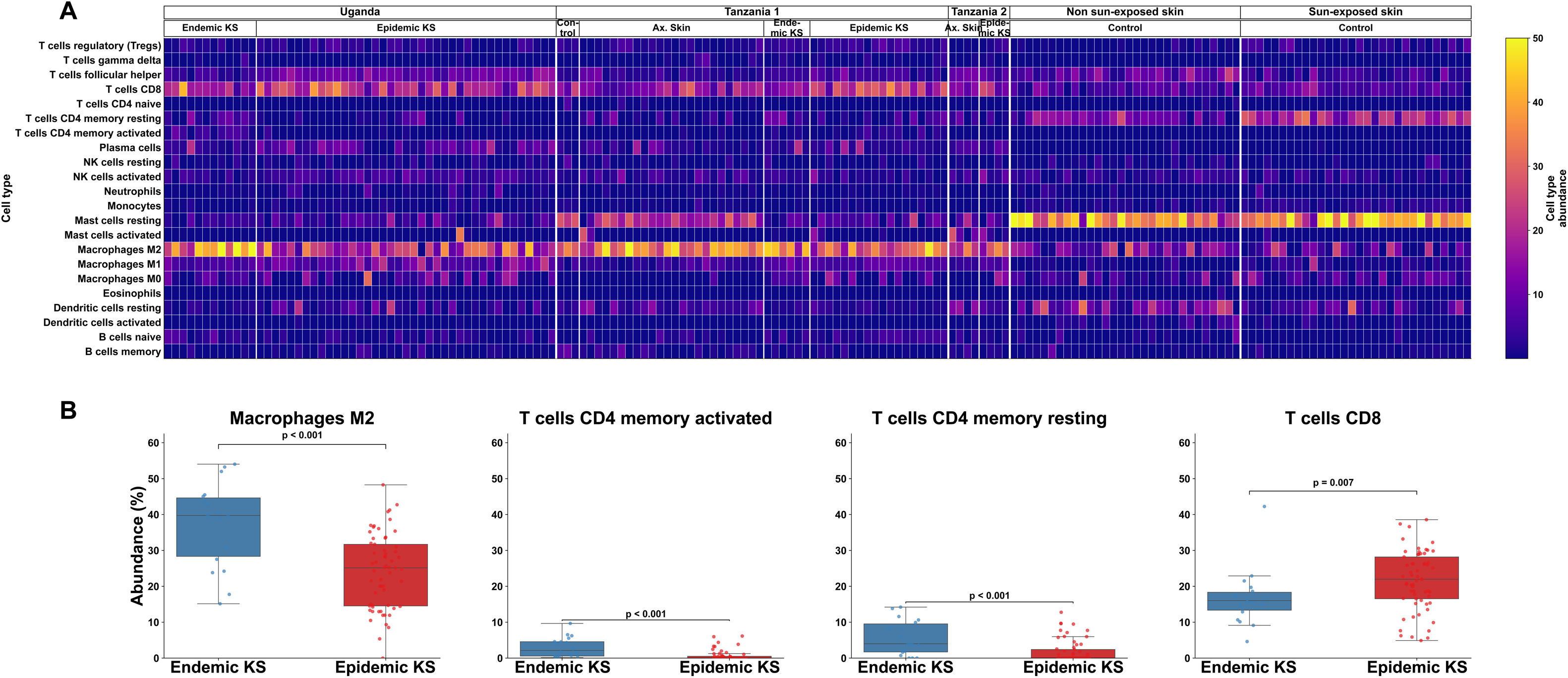
KS tumors show an enrichment of M2 macrophages and CD8^+^ T cells in comparison to control skin samples. (A) Heatmap of relative abundance of the indicated immune cell types found in KS tumors and skin samples from individuals with KS as well as control skin samples from the GTEx consortium, determined via deconvolution of RNA-seq data using CIBERSORTx. The RNA-seq datasets analyzed were derived from (left to right) 12 endemic and 39 epidemic KS tumors from the HIPPOS cohort from Uganda, 3 control skin samples, 24 paired NAT and KS tumor samples from 6 individuals with endemic KS and 18 individuals with epidemic KS from [35]; 4 paired NAT and tumor samples from individuals with epidemic KS from [34]; and 30 non-sun-exposed control skin samples and 30 sun-exposed control skin samples from the GTEx consortium [95]. (B) Plots of the mean and interquartile range (IQR) of the percentages of M2 macrophages, activated CD4**^+^** memory T cells, resting CD4**^+^** memory T cells, and CD8**^+^** T cells, as estimated using CIBERSORTx, in the endemic (blue) and epidemic (red) KS tumors in (A). The upper/lower whiskers extend from the hinge to the largest/smallest value, respectively, no further than 1.5 * IQR from the hinge. Significance between comparisons is indicated by ns (p > 0.05), ** (p<= 0.01), *** (p<= 0.001), **** (p <= 0.0001).

**S3 Fig.**
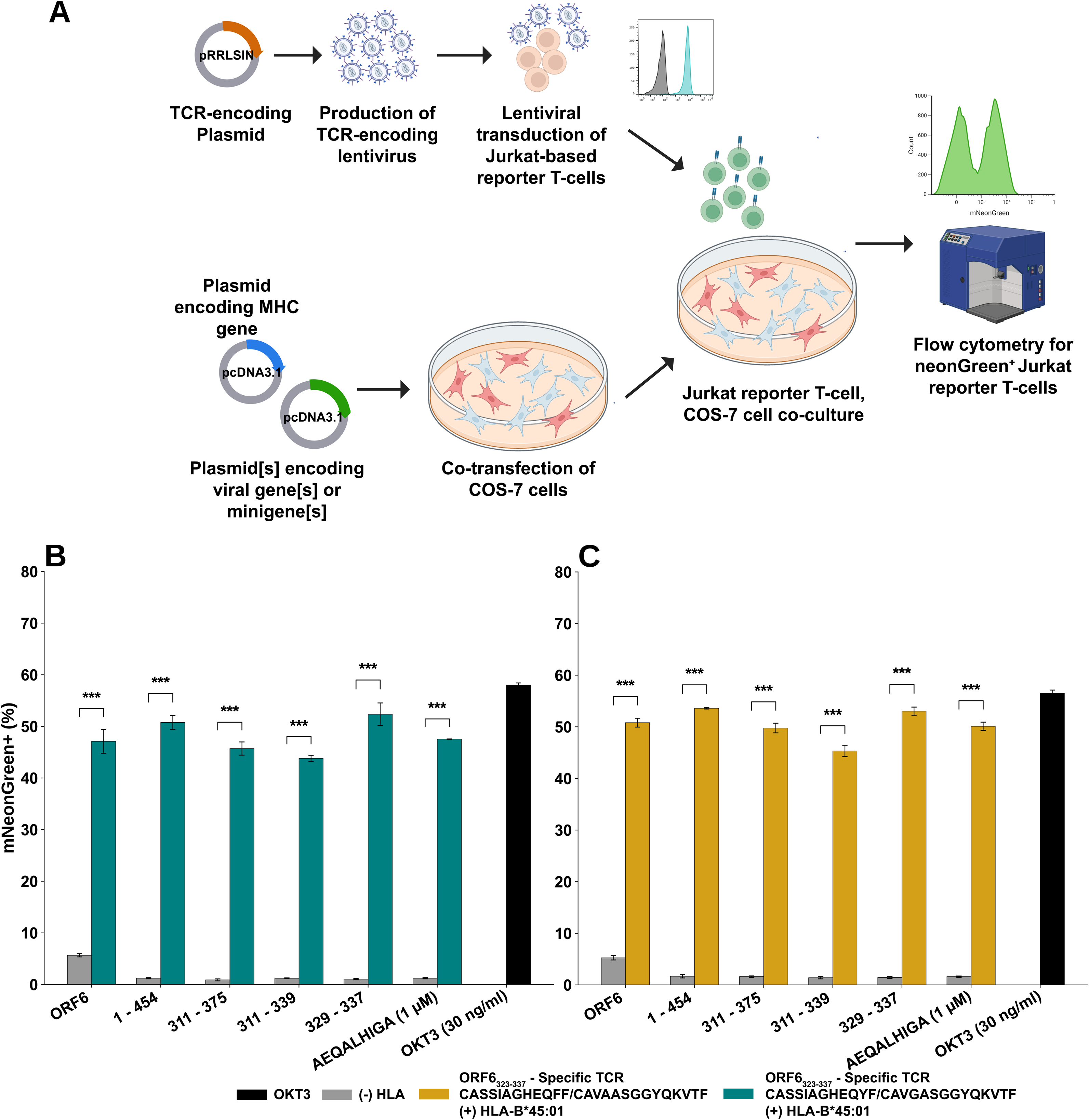
**(A)** Overview of workflow used to define the antigenic specificity of putative KSHV-specific TCRs, utilizing Jurkat reporter T cells transduced with candidate KSHV-specific TCRs and co-cultured with COS-7 cells transiently co-transfected with plasmids encoding the predicted class I MHC restricting allele and KSHV open reading frames. **(B)** Stimulation of TCR-3-expressing Jurkat reporter T cells after co-culture with COS-7 cells transiently transfected with KSHV ORF6, progressively smaller ORF6 minigenes, or the indicated ORF6 peptide. OKT3, positive control. Results for ORF6-specific TCR-2.

**S4 Fig.**
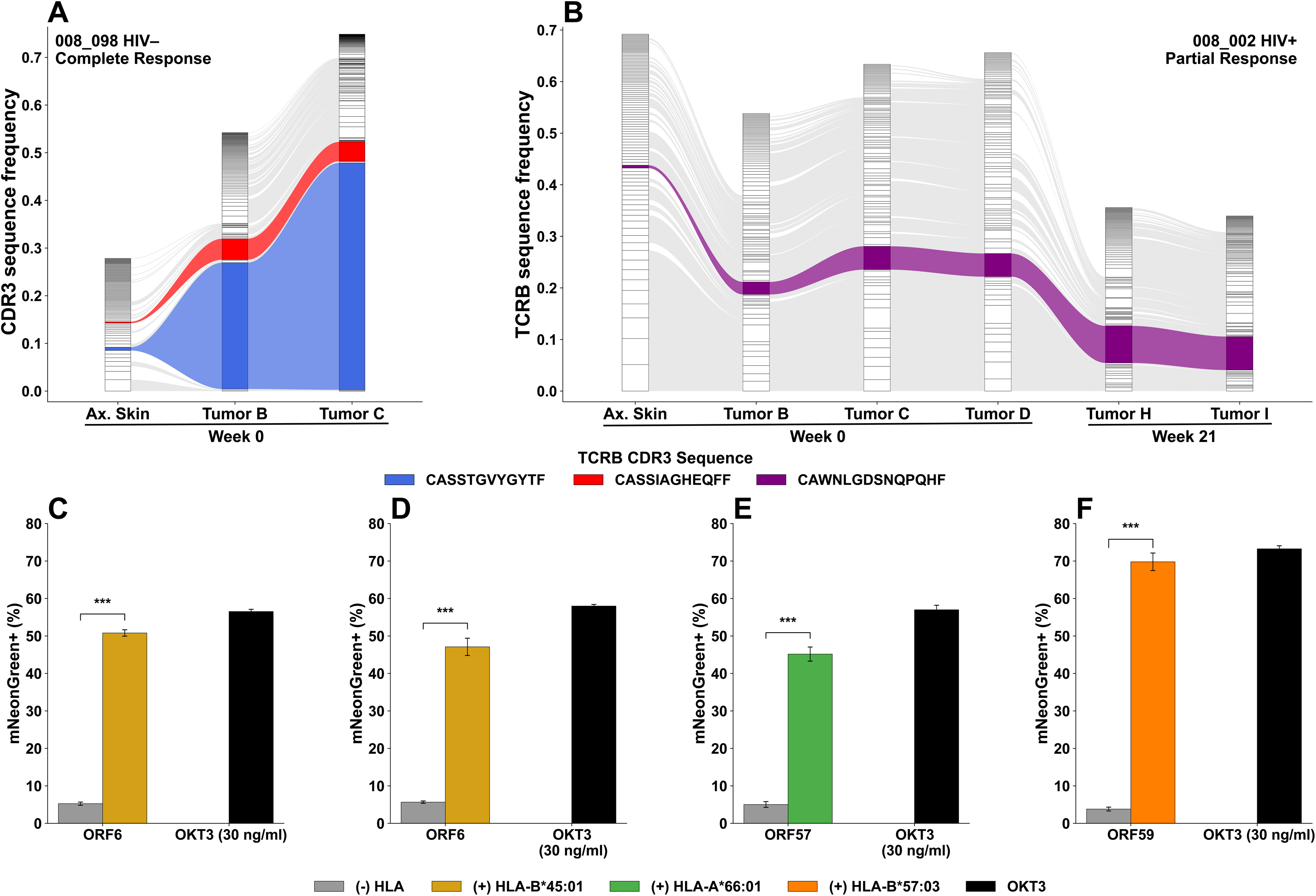
*TRB* CDR3 sequence frequency of KSHV-specific TCRs. **(A)** Alluvial plot showing CDR3 β sequence frequency in biopsies of axillary skin or two different KS lesions in an HIV– individual with KS who achieved a complete remission with chemotherapy. Red indicates the HLA-B*45:01^+^ restricted ORF6-specific TCR-2, and blue indicates the HLA-B*66:01-restricted ORF57-specific TCR-21. **(B**) Alluvial plot showing CDR3 β sequence frequency in biopsies of axillary skin or 5 different KS lesions in an HLA-B*59:01^+^, HIV^+^ individual taken on the initial visit or at week 21 who had a partial response. Purple indicates the HLA-B*57-restricted ORF59-specific TCR. **C-E)** TCR signaling in Jurkat reporter cells (mNeonGreen fluorescence) transduced with the indicated ORF-specific TCRs and co-cultured with LCLs expressing the indicated restricting HLA alleles compared to the response induced by direct CD3 stimulation by OKT3. Mean +/- SEM.

